# Graded functional organisation in the left inferior frontal gyrus: evidence from task-free and task-based functional connectivity

**DOI:** 10.1101/2023.02.02.526818

**Authors:** Veronica Diveica, Michael C. Riedel, Taylor Salo, Angela R. Laird, Rebecca L. Jackson, Richard J. Binney

**Affiliations:** Cognitive Neuroscience Institute, Department of Psychology, School of Human and Behavioural Sciences, Bangor University, Wales, UK; Department of Physics, Florida International University, Miami, FL, USA; Department of Psychiatry, University of Pennsylvania, Philadelphia, PA, USA; Department of Psychology & York Biomedical Research Institute, University of York, UK

**Keywords:** ventrolateral prefrontal cortex, connectivity-based parcellation, resting-state fMRI, meta-analysis, executive function

## Abstract

The left inferior frontal gyrus (LIFG) has been ascribed key roles in numerous cognitive domains, including language, executive function and social cognition. However, its functional organisation, and how the specific areas implicated in these cognitive domains relate to each other, is unclear. Possibilities include that the LIFG underpins a domain-general function or, alternatively, that it is characterized by functional differentiation, which might occur in either a discrete or a graded pattern. The aim of the present study was to explore the topographical organisation of the LIFG using a bimodal data-driven approach. To this end, we extracted functional connectivity (FC) gradients from 1) the resting-state fMRI time-series of 150 participants (77 female), and 2) patterns of co-activation derived meta-analytically from task data across a diverse set of cognitive domains. We then sought to characterize the FC differences driving these gradients with seed-based resting-state FC and meta-analytic co-activation modelling analyses. Both analytic approaches converged on an FC profile that shifted in a graded fashion along two main organisational axes. An anterior-posterior gradient shifted from being preferentially associated with high-level control networks (anterior LIFG) to being more tightly coupled with perceptually-driven networks (posterior). A second dorsal-ventral axis was characterized by higher connectivity with domain-general control networks on one hand (dorsal LIFG), and with the semantic network, on the other (ventral). These results provide novel insights into a graded functional organisation of the LIFG underpinning both task-free and task-constrained mental states, and suggest that the LIFG is an interface between distinct large-scale functional networks.

**Significance statement:** To understand how function varies across the LIFG, we conducted a detailed, bimodal exploration of the spatial transitions in its voxel-wise FC patterns. We provide novel evidence of graded changes along two main organisational axes. Specifically, the LIFG was characterized by an anterior-posterior gradient, which could reflect a shift in function from perceptually-driven processing to task-oriented control processes. Moreover, we revealed a dorsal-ventral shift in FC that is consistent with the idea that domain-specificity is a core principle underpinning functional organisation of the LIFG. These gradients were replicated across task-free and task-constrained FC measures, suggesting that a similar fundamental organisation underpins both mental states.

## 1. Introduction

The left inferior frontal gyrus (LIFG) is ascribed a key role in numerous cognitive domains, including language (Friederici, 2011), semantics (Lambon Ralph et al., 2017), action (Papitto et al., 2020), social cognition (Diveica et al., 2021), and executive function (Fedorenko et al., 2013). The extent of this overlap is remarkable, but what is driving it is unknown. One possibility is that LIFG subserves a singular function which manifests as common activation across domains. Alternatively, detailed exploration of its organisation could reveal subregions with multiple functional specialisations.

Some clues are gleaned from detailed studies of cellular micro-structure and white-matter connectivity that date back to Brodmann (Brodmann, 1909). Cytoarchitecture and ‘fibrillo-architecture’ are proposed to determine a region’s functional characteristics by constraining local processing capabilities and the incoming/outgoing flow of information, respectively (Cloutman & Lambon Ralph, 2012; Passingham et al., 2002). Indeed, these data reveal that the LIFG is far from uniform and, instead, comprises at least three sub-regions with distinct cytoarchitecture (Amunts et al., 1999; Schenker et al., 2008; Wojtasik et al., 2020), neurotransmitter receptor distributions (Amunts et al., 2010), and structural connectivity (Anwander et al., 2007; Klein et al., 2007; Neubert et al., 2014; Wang et al., 2020). However, it has thus far proven difficult to map these structural distinctions onto functional topographies derived from neuroimaging data.

Various functional dissociations have been identified within the LIFG by means of functional neuroimaging, including distinctions between semantic and phonological language processes (Devlin et al., 2003), and between memory retrieval and post-retrieval selection (Badre & Wagner, 2007). However, they have arisen primarily from experimental cognitive approaches and limited neuroimaging datasets which are poorly suited to generating unifying accounts that explain multiple phenomena. A promising alternative is to take a large-scale data-driven approach that spans cognitive domains (Genon et al., 2018). On this basis, one might encapsulate the full functional repertoire of a brain region.

Functional connectivity (FC) patterns derived from neuroimaging data could prove useful because they capture the extent to which regional activation covaries over time and, therefore, are sensitive to context-dependent inter-regional interactions. Moreover, they can reveal aspects of the connectome that might not manifest within other modalities; FC can arise between anatomically remote brain areas without direct structural connections (Damoiseaux & Greicius, 2009; Suárez et al., 2020). The small number of studies that have attempted to divide the LIFG into sub-regions based on FC reveal a heterogenous functional architecture (Clos et al., 2013; Kelly et al., 2010). However, these prior investigations have implemented ‘hard’ clustering algorithms (Eickhoff et al., 2015), which assume that sharp borders separate intrinsically homogeneous neural regions. This means they may fail to identify graded transitions that (a) could give rise to functionally intermediate areas (Bailey & Von Bonin, 1951; Rosa & Tweedale, 2005) and (b) have been observed in the connectivity patterns of other brain regions (e.g., Bajada et al., 2017; Cerliani et al., 2012; Jackson et al., 2018, 2020; Tian & Zalesky, 2018), as well as within the cytoarchitecture of transmodal cortex (Brodmann, 1909). Therefore, the possibility of graded functional differences in the LIFG remains unexplored.

Insights into the nature of spatial transitions in cortical organisation, or *gradients*, can be gleaned using an emergent analytical approach (Bajada et al., 2020; Huntenburg et al., 2018). A key feature of gradient analyses is that they do not presuppose the nature of variation and, therefore, can be used to demonstrate both graded changes and discrete boundaries (Bajada et al., 2017; Jackson et al., 2018; Johansen-Berg et al., 2004). Moreover, they can distinguish between superimposed but orthogonal spatial dimensions of functional variation which might otherwise appear as a singular aspect of organisation (Haak et al., 2018). Despite these advantages, the gradient approach has not yet been applied to studying the connectivity of the LIFG.

Therefore, our aim was to use gradient analyses on FC data in order to 1) elucidate the principal axes of functional organisation within the LIFG and 2) assess whether there is evidence for graded functional differences.

## 2. Methods

We used a data-driven approach to extract LIFG gradients based on two measures of FC: 1) correlations in task-free fMRI time-series and 2) meta-analytically derived patterns of task-driven co-activation from across multiple cognitive domains. This bimodal approach not only allowed us to validate our results using independent datasets, it made it possible to assess the generalisability of the functional organisation of the LIFG across different mental states. Indeed, one data type captures activation patterns associated with spontaneous thought (e.g., a state of mind-wandering; Chou et al., 2017; Doucet et al., 2012), while the other is assumed to reflect mental processes constrained by extrinsic demands (Laird et al., 2013). The summary of our analytical approach is as follows. For each voxel, we 1) extracted BOLD fluctuations over time from resting-state fMRI scans, and 2) meta-analytically identified the brain voxels with which it consistently co-activates across a broad range of task demands. Then, for each FC modality, we compared the fMRI time-series/co-activation patterns of each pair of voxels within the LIFG region of interest (ROI) (see Sections 2.3.1 – 2.3.2). We conducted gradient analyses on the resulting similarity matrices to extract the principal axes of variation and to estimate the degree of gradation (see Section 2.3.3). In a second step, we conducted descriptive analyses to understand which FC differences gave rise to these gradients (see Section 2.3.4). To this end, we performed seed-based resting-state FC and meta-analytic co-activation modelling (MACM) analyses on hard clusters extracted from the extreme ends of the identified gradients. Finally, we probed the functional/task terms (e.g., “cognitive control”, “language”) associated with these IFG sub-regions using functional decoding analyses. A schematic overview of the analytic pipeline is illustrated in Figure 1.

**Figure 1.**
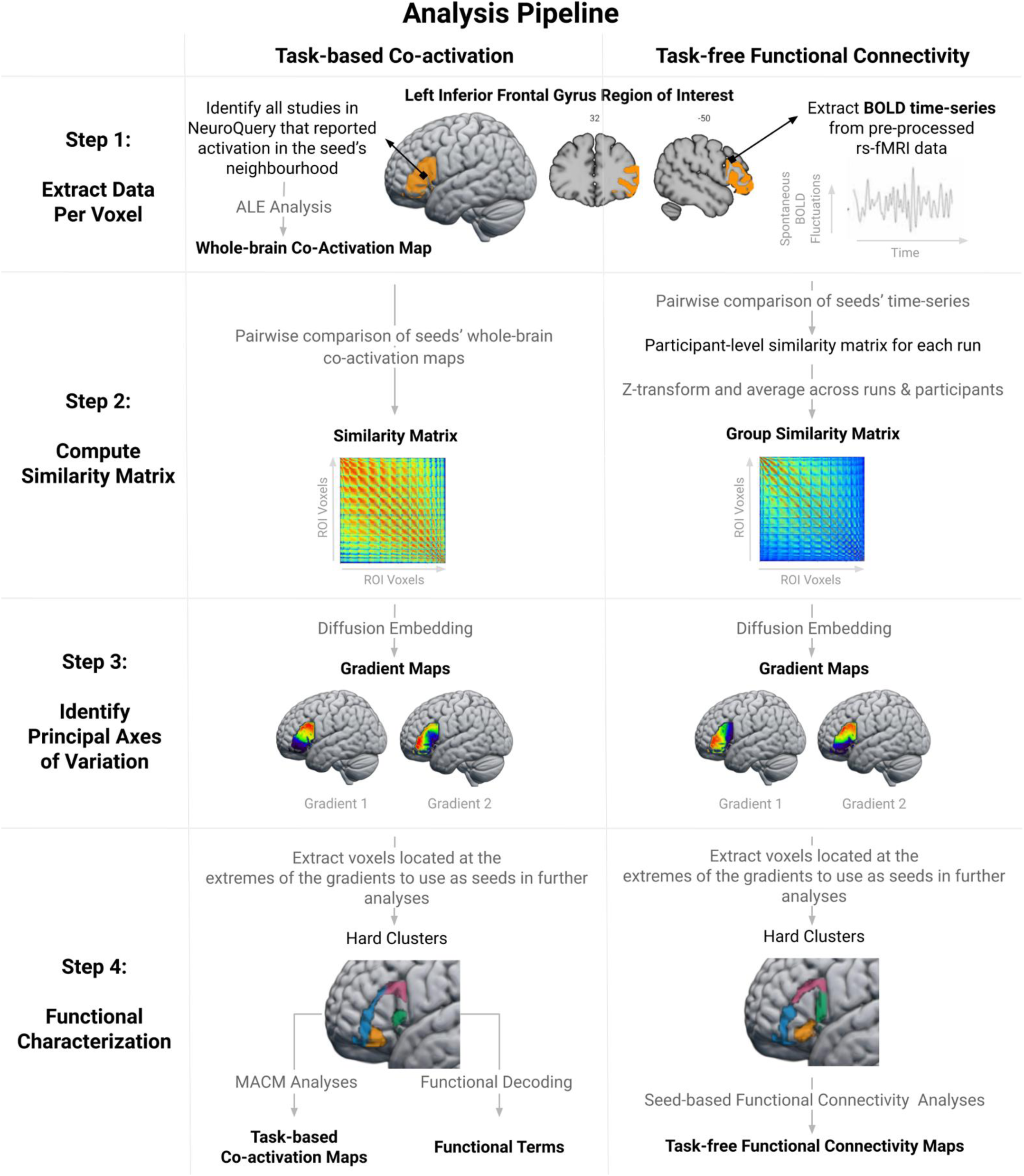
Schematic overview of the analytic pipeline. The output of each analysis step is highlighted in bold. In the first step, we estimated the whole-brain co-activation patterns of individual ROI voxels using meta-analytic co-activation modelling (first column) and extracted their resting-state BOLD time-series (second column). Then, we performed a pairwise comparison of each voxel’s co-activation patterns/time-series using the product-moment correlation coefficient. This resulted in two similarity matrices, which were subsequently used as input for gradient analyses (diffusion embedding algorithm) in order to identify the main axes of variation across the ROI. In the final step, we performed MACM, functional decoding and seed-based resting-state FC analyses on hard clusters extracted from the edges of the gradient maps in order to identify the patterns of FC that characterize different IFG sub-regions. The code used for data analysis can be accessed at: osf.io/u2834/.

### 2.1. Definition of the LIFG region of interest

The LIFG ROI was created by combining the pars opercularis, pars triangularis, and pars orbitalis as delineated in the second release of the Automated Anatomical Labeling (AAL2) atlas (Rolls et al., 2015). In addition, we included the region termed lateral orbital gyrus in the AAL2 parcellation because it is considered to pertain to pars orbitalis (Keller et al., 2009). These regions correspond roughly to Brodmann areas 44, 45 and (part of) 47. We retained only the voxels with 50% or greater probability of being grey matter according to the ICBM-152 template (Fonov et al., 2011). To ensure the ROI did not encompass regions within neighbouring gyri that were of no interest to the present study, the ROI was manually cleaned by removing voxels that crossed gyral boundaries into the precentral gyrus and middle frontal gyrus in the MNI-152 T1 template included in FSL (version 6.0.1). The final ROI comprised 1,813 (2×2 mm) voxels and is depicted in Figure 1 (step 1) and available at: osf.io/u2834/.

### 2.2. Data

#### 2.2.1. Resting-state fMRI data

To assess the functional organisation of the LIFG based on task-free FC, we used the resting-state fMRI time-series of 150 randomly-selected healthy young adult participants (77 females) from the Human Connectome Project S1200 release (Van Essen et al., 2013). For each participant, data were available from up to four 15-minute runs of resting-state fMRI scans collected using the acquisition protocol described by Smith et al. (2013). All four scans were available for 139 participants (92.7% of participant sample), only three scans for three participants (2%) and only two scans for eight participants (5.3%). The data were already pre-processed in MNI space using the minimal processing pipeline described by Glasser et al. (2013) and de-noised using ICA-FIX (Salimi-Khorshidi et al., 2014). We regressed the global signal to further reduce the effects of motion artefacts (Burgess et al., 2016), and smoothed the images using a 4-mm full-width half-maximum Gaussian kernel. In keeping with other resting state studies, we took an additional step of band-pass filtering the data to retain only frequencies between .01 and .08 Hz (Satterthwaite et al., 2013).

#### 2.2.2. Meta-analytic functional neuroimaging data

To assess the functional organisation of the LIFG based on task co-activation patterns, we adopted a meta-analytic approach and capitalized on the openly available NeuroQuery database (neuroquery.org). NeuroQuery contains over 400,000 activation coordinates that were automatically-extracted from 13,459 neuroimaging studies (Dockès et al., 2020). The database also includes estimates of frequency of occurrence of 6308 terms (e.g., ‘cognitive control’, ‘semantic memory’) in each full-text publication from this corpus, which were used to perform functional decoding (see Section 2.3.4).

### 2.3. Data analysis

#### 2.3.1. Task-free FC similarity matrix

To compute the task-free FC similarity matrix, we first extracted the blood oxygen-level-dependent signal time-series of every voxel within the ROI, resulting in a voxel by timepoint matrix for each participant and each run. Then, we computed a cross-correlation matrix by calculating the product-moment correlation coefficient between the time-series of all pairs of ROI voxels. The resulting voxel by voxel matrix was z-score normalized to allow the result of each run to be averaged (Dunlap et al., 2013), in order to generate an average similarity matrix across runs per participant. These participant-level matrices were subsequently averaged resulting in a group similarity matrix. This task-free FC-based similarity matrix was transformed back from z-scores to correlation values for gradient decomposition (see Section 2.3.3).

#### 2.3.2. Task-based co-activation similarity matrix

To compute the task-based co-activation similarity matrix, we first used MACM analyses to identify the brain areas consistently co-activated with each voxel within the ROI. MACM uses meta-analytic data to quantify the co-occurrence of activation between voxels across a broad range of task demands (Laird et al., 2013). This analysis involved extracting all studies in the NeuroQuery database that reported at least one activation peak within 6-mm of a given voxel. Next, we quantified the convergence of activation across the identified experiments using the revised activation likelihood estimation (ALE) algorithm (Eickhoff et al., 2012) as implemented in the Python library NiMARE (Salo et al., 2022). This process was repeated for all voxels within the ROI, resulting in 1,813 unthresholded MACM maps that estimate the strength of co-activation between each ROI voxel and all other brain voxels (ROI voxel by brain voxel matrix). In the second step, we generated a cross-correlation matrix by calculating product-moment correlation coefficient between the MACM map values of each pair of ROI voxels. The resulting task-based co-activation similarity matrix was used as input for the gradient analysis (see Section 2.3.3).

#### 2.3.3. Gradient analysis

We conducted gradient analyses to separately explore the principal axes of variation in task-free FC and task-based co-activation patterns across the ROI. To this end, we first sparsified the similarity matrices by retaining only the top 10% of values row-wise and computed a symmetric affinity matrix using a cosine kernel. The application of this threshold ensures that the results are only based on strong connections, rather than weak and potentially spurious connections (Vos de Wael et al., 2020).

Then, we generated gradient maps by using the diffusion embedding algorithm as implemented in the BrainSpace Python toolbox (Vos de Wael et al., 2020). Diffusion embedding is a type of non-linear dimensionality reduction based on graph theory that describes the high-dimensional connectivity data in terms of distances in a low-dimensional Euclidian space, where the distance between nodes (i.e., voxels) reflects the strength of their connections (i.e. similarity in FC patterns) (for a detailed description, see Coifman & Lafon, 2006). The diffusion embedding algorithm forces voxels with many and/or strong connections closer together and voxels with few and/or weak connections further apart in the embedding space (resulting in gradient maps). We extracted 10 gradients from each modality-specific matrix, but we further interrogate only the first two gradients as they explained considerably more variation in the data compared to the remaining gradients (see Figure S1^1^).

We quantified the degree of gradation in FC changes across the LIFG by estimating the normalised algebraic connectivity of the similarity matrices. This value corresponds to the second largest eigenvalue of the Laplacian of the matrix and represents a descriptive index of how well connected a graph is (Fiedler, 1973). It ranges from zero, which indicates that the graph comprises at least two completely disconnected sub-graphs, to a value of one, which suggests that the graph is characterised solely by graded differences. Thus, the normalised algebraic connectivity of the similarity matrices are indicative of whether the LIFG comprises at least two sharply delineated sub-regions or graded transitions between sub-regions with differences in connectivity/co-activation patterns (Bajada et al., 2020). We note that, while this value is influenced by the smoothing of neuroimaging data, a value much higher than 0 and close to the maximal value possible of 1 is unlikely to be caused only by artificially induced local gradation (Bajada et al., 2017, 2019, 2020; Jackson et al., 2020). We separately estimated the algebraic connectivity of the task-based co-activation matrix and the group task-free FC matrix. In addition, we assessed the gradation in task-free FC matrices at the participant level. This was done to avoid relying only on a gradation metric derived based on the group matrix which is generated by subjecting the individual-level matrices to an additional transformation that may bias the gradation metric.

#### 2.3.4. Functional characterization

While the gradient analysis can estimate the main directions of functional changes, this step alone cannot reveal the qualitative differences in the FC patterns that drive the functional organisation of the LIFG. To describe the task-free FC and task-constrained co-activation patterns of distinct IFG sub-regions, we defined hard clusters based on their locations at the extremes of the gradients by extracting the voxels with the 20% lowest and highest gradient values. The hard clusters are depicted in Figure 1 (step 4) and their MNI coordinates are reported in Table S1. In a graded map, voxels located at the gradient poles should differ most in terms of their FC patterns. Thus, contrasting the task-free FC and task-constrained co-activation characteristics of the clusters located at the extremes of the gradients allows the identification of the patterns that have driven the separation between the clusters in the embedding space. These clusters should not be interpreted as a hard parcellation of the LIFG.

To identify the FC patterns for each cluster, we used the clusters defined based on each of the task-free FC gradient maps as seeds in seed-based resting-state FC analyses. These analyses were performed using the Python package Nilearn (Abraham et al., 2014). For each participant, we used the average resting-state fMRI time-series (concatenated across runs; Cho et al., 2021) of all voxels within each cluster as a regressor in a general linear model predicting the time-series of all grey matter voxels. The resulting cluster FC maps were z-transformed and tested for consistency across participants using a one-sample t-test. In addition, to identify the FC specific to each cluster, which is driving the identification of the gradient, paired-samples t-tests were used to generate contrast maps showing the brain regions with greater FC to one hard cluster than the cluster extracted from the opposite end of the same gradient (anterior vs posterior cluster, dorsal vs ventral cluster). The group-level FC maps were thresholded using a family-wise error (FWE) corrected voxel-height threshold of p < 0.05 and the probabilistic threshold-free cluster enhancement approach as implemented in the R package pTFCE (Spisák et al., 2019). We wanted to identify the brain regions that (i) displayed greater functional coupling with one LIFG cluster than the cluster at the opposite end of the same gradient and, at the same time, (ii) were significantly coupled with the respective LIFG cluster. Therefore, the contrast maps (determined using the paired-samples t-tests) were masked by the significant connectivity of each cluster (determined using the one-sample t-tests).

To identify the co-activation patterns of each LIFG cluster, we conducted MACM analyses on seeds defined based on each task-based gradient map using the Python package NiMARE (Salo et al., 2022). Specifically, we ran ALE analyses on all studies from the NeuroQuery database that reported at least one activation peak within the seed (see Table S2 for the number of studies identified for each cluster) to identify the brain regions consistently involved in the studies that activate the seed. Specifically, the resulting MACM maps quantify the convergence of activation across all studies that reported activation within the seed. These maps were thresholded using a FWE corrected voxel-level threshold of p < 0.05. Then we conducted contrast analyses to identify the brain regions that co-activate more consistently with one hard cluster than the cluster extracted from the opposite end of the same gradient (anterior vs. posterior cluster, dorsal vs. ventral cluster). The contrast maps were thresholded using an uncorrected p < 0.05 threshold. To understand which brain regions display (i) greater co-activation with the cluster located at one extreme of the gradient than the other extreme and (ii) significant co-activation with the cluster, we masked the contrast maps by the significant cluster-specific MACM map (determined using independent ALE analysis).

It is important to note that we conducted contrast analyses using the same FC modality (i.e., MACM of clusters extracted from the task-based gradients, seed-based resting-state FC analyses of clusters extracted from the task-free gradients) in order to visualize the differences that have driven the gradients, and not to test whether there were significant FC differences between the clusters. The non-inferential and descriptive nature of these follow-up analyses circumvents analytic circularity (Eickhoff et al., 2015). Nonetheless, we repeated these sets of analyses using an independent FC modality (i.e., MACM of clusters extracted from the task-free gradients, seed-based resting-state FC analyses of clusters extracted from the task-based gradients) to confirm whether the FC maps are consistent regardless of the approach adopted to define the clusters. These analyses revealed similar FC patterns and are only reported in supplementary Figures S8-9.

In line with recent recommendations (Uddin et al., 2022), we determined the network affiliations of our novel findings by comparing them with a commonly-used parcellation scheme. We used the 7-network parcellation proposed by Yeo et al. (2011) as the reference atlas. For each task-free FC and task-based co-activation map, we computed the percentage of voxels that overlap with each of the seven reference networks. The LIFG has been consistently implicated in semantic processing (Jackson, 2021) which is thought to be supported by a functional network that is dissociable from other canonical networks such as the core default network (DN) (Branzi et al., 2020; Humphreys et al., 2015; Jackson et al., 2016, 2019; Jung & Lambon Ralph, 2022). Therefore, we also computed the overlap between our results and a mask of the semantic network (SN) proposed by Jackson et al. (2016). The reference SN map represents the set of regions that were significantly functionally coupled with the left ventrolateral anterior temporal lobe (ATL), which has been attributed a crucial role in semantic cognition (Binney et al., 2010; Jackson et al., 2016; Lambon Ralph et al., 2017). It is of note that we are not able to dissociate between the DN and SN in these analyses (as has been done by, for example, Humphreys et al., 2015; Jackson et al., 2019) because there is a considerable degree of spatial overlap between the DN mask from Yeo et al. (2011) and the semantic network obtained by Jackson et al. (2016).

Lastly, to identify functional terms associated with each cluster as an index of its potential function, we conducted functional decoding analyses using the BrainMap chi-square approach as implemented in NiMARE (Salo et al., 2022). For each term in the NeuroQuery database, the consistency analysis (also known as forward inference) computes the likelihood of activation reported within the seed given presence of the term in the article’s text, whereas the specificity analysis (also known as reverse inference) estimates the posterior probability of an article containing the term given activation reported inside the seed. The results of these analyses were thresholded at p < 0.05 using the Benjamini-Hochberg false discovery rate correction. To aid the interpretability of the results, we retained only the terms with at least 80% likelihood of being related to cognitive functions based on raters’ annotations (Bottenhorn et al., 2019).

## 3. Results

### 3.1. Gradient maps

The first two task-free FC gradients were selected for further analysis because together they accounted for > 50% of variance, while the lower-order gradients explained less than 11% of variance each (Figure S1). The voxels’ gradient values, which reflect the similarity between their resting-state fMRI time-series, were visually coded and projected on the brain using a colour spectrum from red to dark blue to reveal the pattern of change in task-free FC across the LIFG. As can be seen in Figure 2A, the FC patterns of the LIFG are principally organized along an anterior-posterior axis that accounted for 30% of the variance. This gradient progressed from the anterior portion of the LIFG, bordering the inferior part of the inferior frontal sulcus (IFS), to the posterior region, bordering the precentral gyrus. The second gradient, which explained 25% of the variance, revealed changes in connectivity along the superior-inferior dimension. This gradient progressed from the superior part of the IFS and the precentral sulcus to the inferior portion of the IFG, bordering the lateral orbital sulcus. The algebraic connectivity of the group similarity matrix was 0.71, suggesting a high level of gradation in task-free FC changes across the LIFG. This was confirmed by the distribution of the algebraic connectivity of the individual-level similarity matrices (Figure S2) which had a mean of 0.89 (SD = 0.02). The group similarity matrix, reordered based on the voxels’ positions along the first and second gradients, and showing the graded change in FC across voxels in the LIFG, is illustrated in Figure S3.

**Figure 2.**
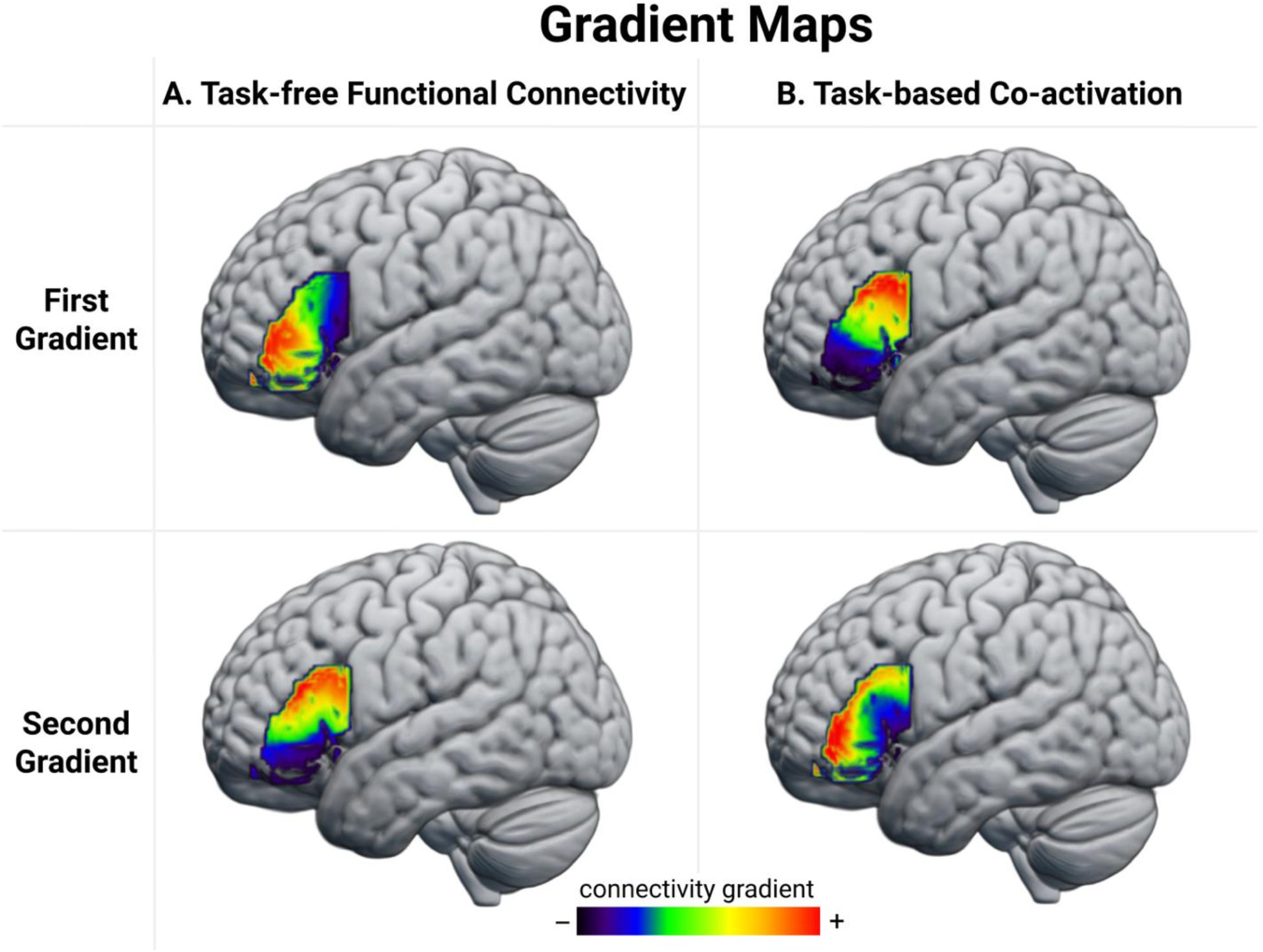
The first two gradient maps extracted from the A) task-free FC similarity matrix and B) task-based co-activation similarity matrix. Compared to regions represented with colours further apart on the colour spectrum, regions represented using colours that are closer together show greater similarity in their A) correlation with each other over time during resting fMRI scans and B) their patterns of co-activation across tasks spanning a range of cognitive domains. The +/- indicate different poles of these gradient dimensions, but the assignment to a specific end of a dimension is arbitrary. The gradient maps can be accessed at: neurovault.org.

The first two task-based co-activation gradients were selected for further analysis because together they accounted for > 60% of the variance, while the lower-order gradients individually explained less than 11% of the variance (Figure S1). The principal gradient accounted for 42% of the variance and progressed along a dorsal-ventral axis from the inferior frontal junction (IFJ) to the antero-ventral region bordering the lateral orbital sulcus and inferior portion of the IFS. The second gradient explained 21% of the variance and revealed changes in connectivity that followed the rostral-caudal axis in a radial pattern progressing from the inferior portion of the pars opercularis towards the IFS. The algebraic connectivity of the co-activation similarity matrix was 0.77, suggesting that LIFG is characterized by gradual changes in consistent patterns of co-activation across cognitive domains (see Figure S3 for the reordered matrices). Because the unit of the task-based analysis is the study rather than the participant, the gradation cannot be assessed at the participant level as in the case of the task-free analysis reported above.

The gradients extracted from the two independent datasets converge on two principal organisational axes of the LIFG: anterior-posterior and dorsal-ventral. Visual inspection of the gradient maps suggests that the first task-free gradient and the second task-based gradient capture a similar anterior-posterior axis of functional variation, which is supported by a strong positive correlation of 0.77 between voxels’ position ranks on the two gradients (see Figure S4 for the scatterplot). Likewise, the second task-free gradient and the first task-state gradient capture a similar dorsal-ventral organisational dimension. This observation is supported by a strong positive correlation of 0.7 between voxels’ position ranks on the two gradients (Figure S4). The orders in which these gradients appear are switched between the task-free and task-constrained FC data, and this is because of a difference in the relative amount of variance explained by each gradient. Because it is subtle relative to the similarities, this difference could be attributable to noise but it may also reflect meaningful differences in the connectivity revealed by task-free and task-constrained mental states (Eickhoff & Grefkes, 2011).

### 3.2. Functional characterization

#### 3.2.1. Differential task-free FC patterns

We contrasted the whole-brain resting-state connectivity patterns of the clusters located at the extremes of the anterior-posterior task-free gradient. This revealed differences in their functional coupling with a bilateral and distributed set of brain regions (Figure 3A; Table S3). The anterior cluster showed stronger FC with frontal regions, including the right IFG (pars orbitalis), bilateral IFS, dorsal and orbital portion of the middle frontal gyrus (MFG), superior frontal gyrus (SFG), medial prefrontal cortex (mPFC), and orbitofrontal cortex (OFC), with parietal regions in the posterior cingulate cortex (PCC), angular gyrus (AG), and inferior parietal lobule (IPL), and with temporal regions along the length of the middle and inferior temporal gyri (MTG/ITG) and in the fusiform gyrus (FG), and left hippocampus. In contrast, the posterior cluster showed stronger FC with frontal regions in the right IFG (pars opercularis), the middle portion of the MFG, precentral gyrus, pre-supplementary motor area (pre-SMA), anterior and middle cingulate cortex (ACC), and insula, and with posterior cortical regions in the supramarginal gyrus (SMG), and posterior superior temporal sulcus, and with basal ganglia.

**Figure 3.**
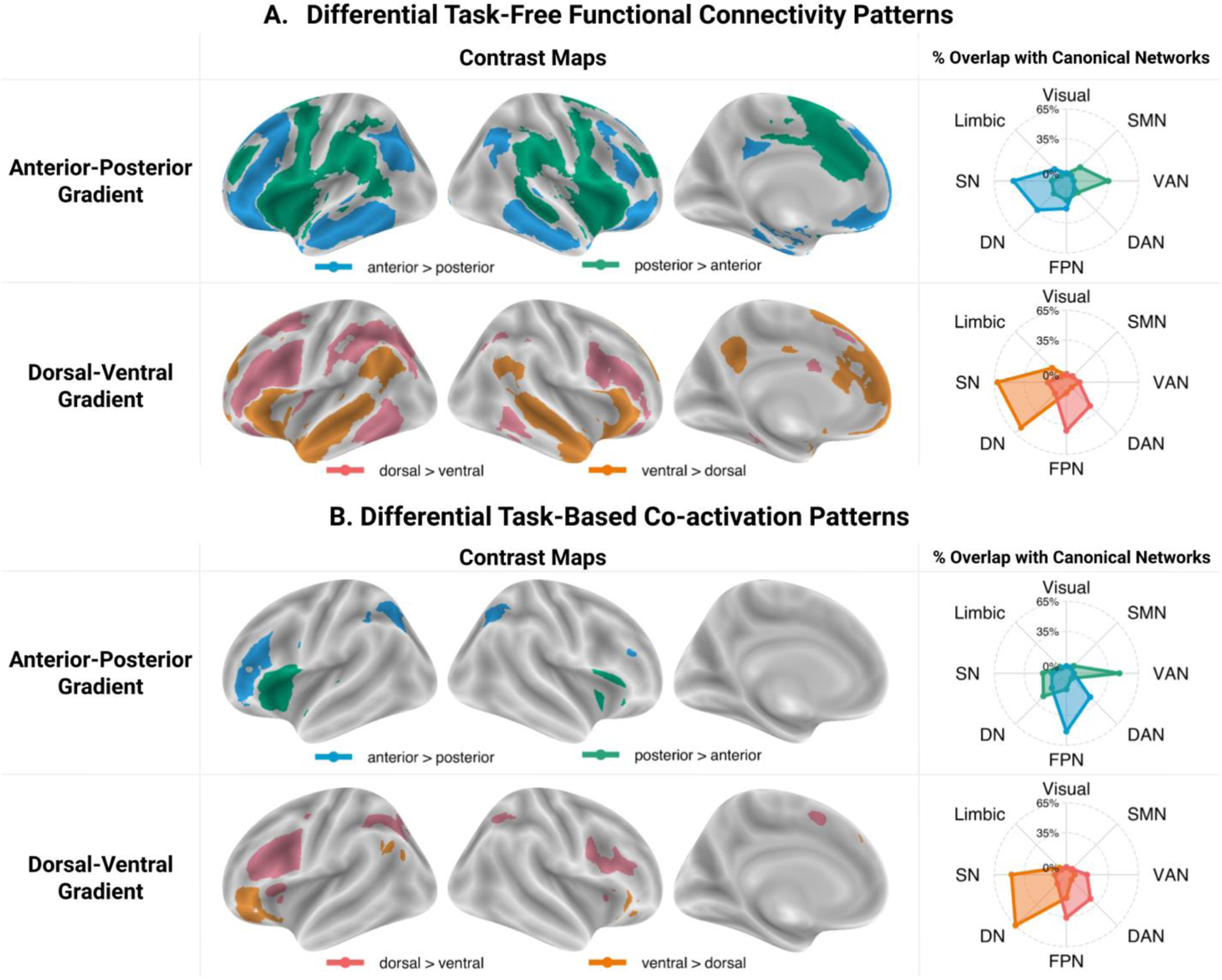
Results of contrast analyses between A) the task-free connectivity patterns (derived using seed-based resting-state FC analyses) of the IFG clusters located at the extremes of the anterior-posterior and dorsal-ventral task-free gradients and B) task-constrained co-activation patterns (derived using MACM analyses) of the IFG clusters located at the extremes of the anterior-posterior and dorsal-ventral task-based gradients. These contrast maps were masked using cluster-independent maps. The spider plots in the right column show the percentage of voxels in each contrast map that overlap between with each of the seven canonical networks from Yeo et al. (2011), as well as the SN from Jackson et al. (2016) (available at: github.com/JacksonBecky/templates). Note that percentage values are relative to the size of each contrast map; therefore, only the relative patterns of overlaps within each contrast map are of interest and direct comparisons between the network affiliations of different contrast maps are misleading. The contrast maps can be accessed at: neurovault.org/collections/ETPEFCDV/. Abbreviations: SMN - somatomotor network, VAN – bentral attention network; DAN – dorsal attention network; FPN – frontopariental control network; DN – default network; SN – semantic network.

Comparison between the task-free FC of the dorsal and ventral clusters revealed stronger coupling between the dorsal LIFG and frontal regions in bilateral IFS and IFJ, left MFG, and pre-SMA, parietal cortex in bilateral IPS, and left IPL, and temporal cortex in bilateral posterior ITG and left FG (Figure 3A; Table S4). In contrast, the ventral LIFG showed increased connectivity to the frontal cortex in the right IFG (pars triangularis and pars orbitalis), bilateral SFG, mPFC, and ACC, to the precuneus, a swathe of temporal cortex progressing from the bilateral ventrolateral ATL through the MTG towards the AG, and to the left hippocampus.

Comparison between the cluster-specific task-free FC patterns and canonical networks indicate stronger functional coupling between the anterior LIFG and regions falling within the bounds of the SN/DN and the frontoparietal network (FPN), and between the posterior LIFG and brain regions associated with the ventral attention network (VAN) and somatomotor network (SMN). The dorsal LIFG showed stronger FC with regions of the FPN and dorsal attention network (DAN), whereas the ventral LIFG showed a preference for SN/DN regions. Additional conjunction analyses showed that both the anterior and posterior clusters are coupled with regions of the FPN and SN/DN, and that both the dorsal and ventral clusters are functionally connected mainly with SN/DN regions (Figure S7A; Table S3-4).

#### 3.2.2. Differential task-constrained co-activation patterns

The anterior LIFG showed increased consistent co-activation across a wide variety of tasks with frontal regions in the right IFS and precentral gyrus, and also with bilateral IPS and left posterior ITG, whereas the posterior LIFG co-activated more with the right IFG, bilateral anterior insula and left superior temporal gyrus (Figure 3B; Table S5). The dorsal cluster co-activated more with frontal cortex in the right IFJ, bilateral precentral gyrus, dorsal anterior insula, and pre-SMA, and with the IPS, and left posterior FG (Figure 3B; Table S6). In comparison, the ventral cluster showed increased co-activation with the right IFG (pars orbitalis), and left mPFC, MTG and AG. Given the conservative threshold applied to the independent maps, we also looked at the whole contrast maps without masking by these independent maps. These additionally revealed more consistent co-activation of the posterior cluster with the bilateral STG and of the ventral cluster with the bilateral ATL, precuneus and left AG (Figure S6).

Comparison between the cluster-specific task-based co-activation patterns and canonical networks shows that the anterior LIFG cluster co-activates more consistently with brain regions that are part of the FPN and DAN, whereas the posterior LIFG cluster co-activates mainly with regions associated with the VAN and, when additional masking is not applied, the SMN. The dorsal cluster co-activates preferentially with regions of the FPN and DAN, whereas the ventral LIFG cluster shows stronger co-activation with the DN/SN. Additional conjunction analyses showed overlap between the co-activation maps of the anterior and posterior clusters and those of dorsal and ventral clusters primarily in regions of the FPN (Figure S7B; Table S5-6).

The FC analyses performed on clusters extracted from the gradient maps derived using the independent dataset (i.e., seed-based FC analyses of clusters derived using NeuroQuery studies and MACM analyses of clusters derived using task-free fMRI time-series), which were conducted to assess the robustness of the results across different strategies for defining seeds, revealed a similar pattern of results (Figure S8-9).

#### 3.2.3. Comparison between the task-free and task-based FC patterns

In sum, the task-free and task-based analyses implicate overlapping regions, although the clusters identified in the task-based analyses were less extensive. Specifically, the anterior LIFG was connected with executive control regions (e.g., IFJ, IPS; Assem et al., 2020; Camilleri et al., 2018; Fedorenko et al., 2013), but in the task-free maps it was also connected to regions implicated in semantic cognition (e.g., ATL, AG; Binder & Desai, 2011; Lambon Ralph et al., 2017). Further, the posterior LIFG was connected to areas that have been ascribed important roles in sensorimotor processing, as well as in phonological and articulatory linguistic processes (e.g., bilateral STS/STG, but in the task-free maps it was also connected to motor and premotor cortices, SMA, MFG and SMG; Hartwigsen et al., 2010; Hickok, 2009; Hickok & Poeppel, 2007; Price, 2012; Ueno et al., 2011; Vigneau et al., 2006). This cluster was also connected to regions considered crucial for salience processing (e.g., anterior insula, but in the task-free results also to dorsal ACC; Menon & Uddin, 2010; Uddin, 2015). The dorsal LIFG was connected to regions that are implicated in executive function (e.g., IFJ, MFG, IPS; Assem et al., 2020; Camilleri et al., 2018; Fedorenko et al., 2013), whereas the ventral LIFG was connected with a set of regions ascribed key roles in semantic and episodic memory (e.g., ATL, medial temporal lobe, AG; Binder & Desai, 2011; Lambon Ralph et al., 2017).

Despite the similarities in the regions implicated, there were some differences in the network affiliations derived from the task-free and task-based analyses. However, comparing the network affiliations of the different contrast maps directly is not possible because 1) the overlap index depends on the size of the maps, which differs considerably between the task-free and task-based analyses and 2) there are differences between the seeds upon which the task-free and task-based analyses are based (see Figure 1;e.g., the task-based anterior seed extends across the length of the IFS and overlaps with the dorsal LIFG seed, whereas the anterior seed used for the task-free analysis does not). Therefore, we will focus the interpretation on the similarities.

The dorsal LIFG connected to FPN and DAN regions, two networks that contribute to the task-general multiple demand network (MDN; Assem et al., 2020; Majerus et al., 2018). In contrast, the ventral LIFG was affiliated mainly with the DN/SN. The DN and SN cannot be distinguished in our assessment given the high degree of spatial overlap between the masks used. However, we note that both the task-free and the unmasked task-based results suggest strong coupling with the ATL, a key hub of semantic knowledge (Lambon Ralph et al., 2017), as well as with the left hippocampus/parahippocapal gyrus, known to be important for episodic memory (Burgess et al., 2002; Dickerson & Eichenbaum, 2010). As such, the dorsal-ventral organisational dimension seems to distinguish between domain-general control networks at the dorsal end and memory-related networks at the ventral end.

The posterior LIFG showed a preference for the VAN, suggestive of a role in perceptually-driven cognition (Corbetta et al., 2008; Corbetta & Shulman, 2002). The anterior LIFG showed a preference for regions that overlap with the FPN, consistent with a role in cognitive control (Assem et al., 2020). The task-free data revealed additional strong coupling with regions that are part of the DN/SN. The task-based analyses might have led to less extensive association with the DN because this network is known for its tendency to deactivate in response to various task demands (Buckner et al., 2005; Mazoyer et al., 2001; Shulman et al., 1997), but it tends to activate during mind wandering states which frequently occur during resting-state scans (Smallwood et al., 2021). Nonetheless, there is evidence that the DN works with the FPN in support of some types of goal-directed cognition (Spreng et al., 2010, 2014) and that it contributes to cognitive control (Crittenden et al., 2015). As such, the anterior-posterior organisational dimension seems to distinguish between higher-order transmodal networks at the anterior edge and perceptually-driven networks at the posterior edge.

#### 3.2.3. Functional decoding

The functional decoding results suggest possible functional associations of the different LIFG clusters. All four LIFG clusters were significantly associated with terms related to semantic and linguistic processing, including *language, semantic, retrieval, reading, and phonological*. Compared to the posterior cluster, the anterior cluster was associated with the terms *executive (function)* and *memory retrieval.* In contrast, the posterior cluster was associated with terms related to perceptual and motor processing, such as *movement, recognition, auditory,* as well as speech-related terms, such as *phonetic* and *vocal.* Compared to the ventral cluster, the dorsal cluster was associated with terms related to a wide range of cognitive/behavioural domains and input modalities, including *visual, auditory, visuospatial, working memory, executive, social, reward, mood.* In contrast, the ventral cluster was associated with terms such as *memories, mentalizing, reappraisal and autobiographical,* which are suggestive of the purported internally-oriented functions of the DN (Smallwood et al., 2021). While functional decoding approaches can provide pointers to the potential task associations of these regions, it is important to note that the specificity of the results is limited due to the limitations of automated data mining tools like NeuroQuery, such as the aggregation of all contrasts reported in an article, regardless of the cognitive aspects they isolate (Dockès et al., 2020). As a consequence, interpretation should focus on the overall patterns that emerge, rather than the associations of individual terms. Detailed lists of the functional associations are presented separately for forward and reverse inference analyses and task-free and task-based clusters in Supplementary Figures S10-11.

Figure 4 summarizes the functional decoding results that were consistent for the clusters extracted from the task-free and task-based gradients (e.g., terms associated with both the anterior edge of the task-free gradient and the anterior extreme of the task-based gradient). It also includes a schematic of the proposed functional organisation, which takes into account the results of the FC contrast analyses, the network affiliations, the functional decoding, as well as previous literature reviewed in detail in the Discussion.

**Figure 4.**
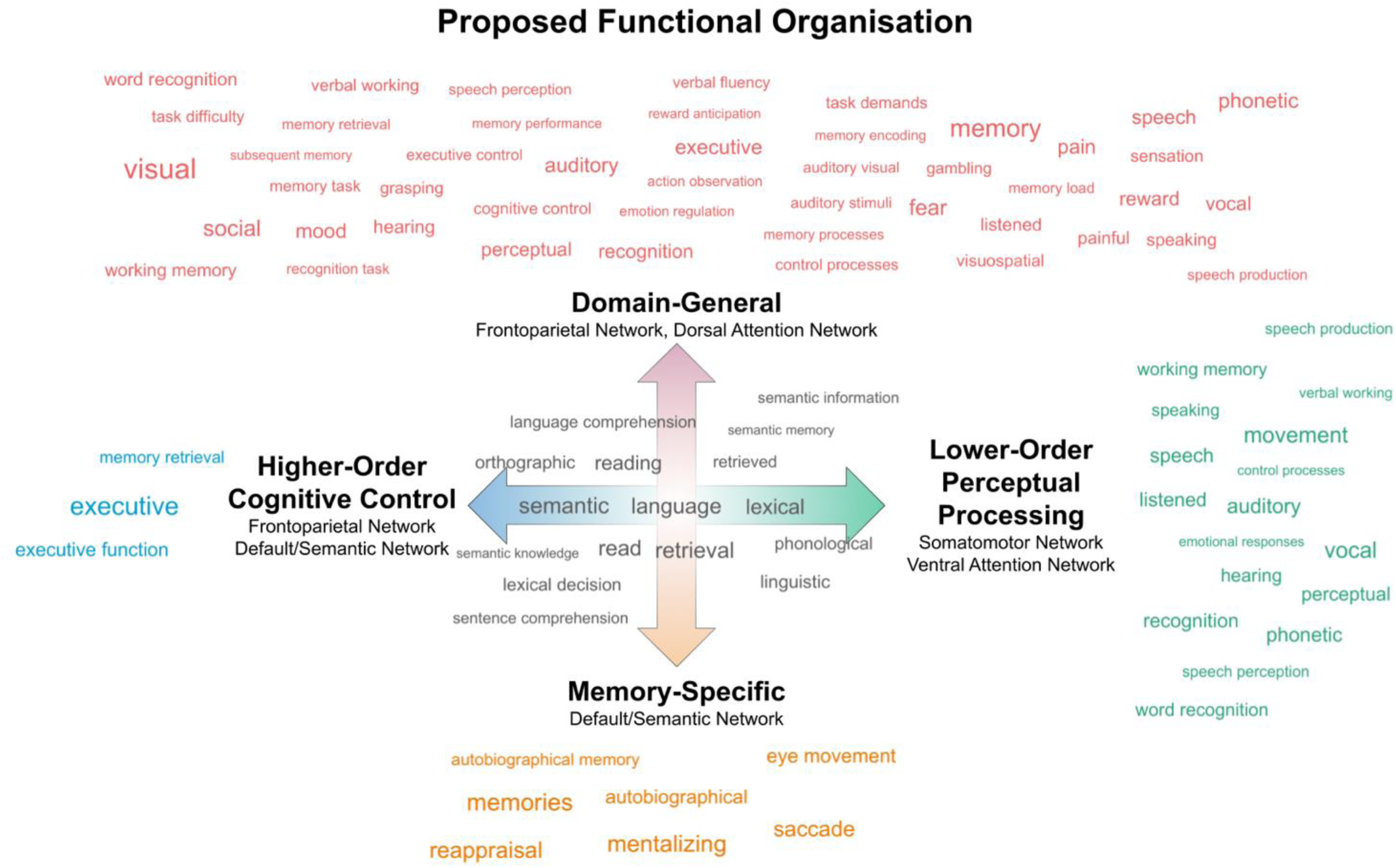
Schematic representation of the proposed functional organisation of the LIFG. The anterior-posterior organisational axis (represented by the horizontal blue-to-green arrow) might reflect a shift from lower-order perceptual processing (posterior LIFG) via affiliation with the SMN and VAN, and higher-order cognitive control (anterior LIFG) via affiliation with the FPN and DN. The dorsal-ventral axis (represented by the vertical orange-to-red arrow) might reflect a shift from domain-general executive functions (dorsal LIFG) via affiliation with the FPN and DAN and the cognitive control of information stored in long-term memory (ventral LIFG) via affiliation with the DN/SN. The word clouds illustrate functional terms associated with the LIFG clusters located at the extremes of the task-free and task-based gradients in the forward or reverse inference functional decoding analyses - terms associated with the anterior but not posterior cluster (blue); the posterior but not anterior cluster (green); the dorsal but not ventral cluster (red); the ventral but not dorsal cluster (orange); and with all four clusters (grey). The size of the word reflects the effect size of the association.

## 4. Discussion

The present study made the first attempt to use data-driven gradient analyses of FC data to elucidate the functional organization of the LIFG. We specifically aimed to 1) map the principal axes of change in function, and 2) determine whether these shifts might be graded. In the following two sections, we shall summarise our novel findings, and then discuss their functional significance.

### 4.1. Graded topographical organisation of the LIFG along two principal axes

Our analyses converged upon two key findings. First, the FC across the LIFG is principally organized along two orthogonal axes. One of these axes is oriented in an anterior to posterior direction and driven by stronger coupling with the FPN and DN in the rostral aspect, and with the VAN and SMN at the caudal end. The second arose along a ventral to dorsal orientation, and reflected greater connectivity of ventral LIFG to the DN, whereas dorsal regions abutting the IFS/IFJ were more tightly coupled with the FPN and DAN. These differential patterns of FC are in line with previous investigations (Barredo et al., 2016; Clos et al., 2013; Davey et al., 2016; Jakobsen et al., 2016, 2018; Kelly et al., 2010; Muhle-Karbe et al., 2016; Neubert et al., 2014; Wang et al., 2020) and suggest that the LIFG interfaces between distinct large-scale functional networks, consistent with its proposed role as a cortical hub (Buckner et al., 2009; Sepulcre et al., 2012).

Our second key finding is that FC of the LIFG shifts in a relatively graded manner. Precisely, the algebraic connectivity of the similarity matrices revealed that FC analyses do not support there being abrupt boundaries and discrete functional parcels in LIFG. This is consistent with both contemporary descriptions of LIFG connectivity based on intraoperative cortico-cortical evoked potentials (Nakae et al., 2020) and structural properties (Binney et al., 2012; Thiebaut de Schotten et al., 2017), as well as classical descriptions that include a fan-shaped set of anatomical projections emanating from the IFG into the lateral temporal lobe (Dejerine & Dejerine-Klumpke, 1895). The present study is the first to confirm a graded organisation of the LIFG using a bimodal dataset, taking into account task-driven variation on one hand, and task-free FC on the other.

Overall, our findings are compatible with previous parcellations despite key differences in the methodological approaches (Clos et al., 2013; Kelly et al., 2010; Klein et al., 2007; Neubert et al., 2014; Wang et al., 2020). This includes those that have taken a ‘hard’ clustering approach to parcellating left prefrontal cortex (PFC). For example, Neubert et al. (2014) parcellated PFC based on structural connectivity and found that the LIFG fractionated into discrete subdivisions positioned along the anterior-posterior dimension. The connectivity of these parcels was distinct from those situated dorsally in the adjacent IFS and IFJ, which implies a further dorsal-ventral dimension of organisation. The co-existence of these two axes of LIFG organisation is also apparent in hard parcellations of the LIFG (Clos et al., 2013; Wang et al., 2020), its right hemisphere homologue (Hartwigsen et al., 2019), and more encompassing parcellations of cortex (Glasser et al., 2016). Of course, the results of graded and hard parcellation are not identical as hard parcellations: (1) force voxels that are part of intermediate regions with gradually-changing connectivity to be within the borders of discrete clusters (Bajada et al., 2017; Haueis, 2012) and (2) require the a priori specification of the number of clusters that are to be identified, perhaps making them insensitive to finer details. However, the two superimposed yet orthogonal modes of organization identified here may have driven prior hard parcellations of the LIFG.

### 4.2. The putative functional significance of the LIFG’s functional connectivity gradients

Taken together, the cluster-specific FC patterns and functional decoding results paint a coherent picture regarding the functional significance of the graded connectivity patterns that appear across the LIFG. On this basis, and in conjunction with the results of previous functional neuroimaging studies, we propose the following interpretation, which has also been illustrated schematically in Figure 4. First, the dorsal-ventral axis might reflect a functional transition from domain-general executive function (dorsal LIFG) to domain-specific control of meaning-related representations (ventral LIFG). Second, the anterior-posterior axis might reflect a shift from perceptually-driven processes (posterior LIFG) to higher-level transmodal control (anterior LIFG). We discuss this proposal in further detail below.

The dorsal LIFG was functionally coupled with regions that comprise the FPN and DAN. These two networks contribute to a wide variety of task demands that span multiple cognitive domains (Assem et al., 2020; Cole et al., 2013). In contrast, the ventral LIFG was preferentially affiliated with the DN, as well as brain regions that have been ascribed key roles in semantic cognition, such as the anterior temporal lobes (Binney et al., 2010; Lambon Ralph et al., 2017). Thus, the shift in FC towards ventral IFG subregions might reflect a specialization towards the application of cognitive control to prior knowledge. Indeed, it has been proposed that the LIFG, as a whole, sits in a unique position at the intersection of the MDN and the DN, and that this makes it ideally suited for implementing demanding operations on meaning-related representations (Chiou et al., 2022; Davey et al., 2016). Consistent with this, the LIFG responds reliably to an increased need for the control of semantic information across a wide range of experimental paradigms (Diveica et al., 2021; Jackson, 2021), including those requiring episodic memory retrieval (Vatansever et al., 2021). However, it is increasingly apparent that there are finer-grained functional distinctions within the LIFG; dorsal LIFG regions near IFS/IFJ overlap with the MDN and are engaged by control demands that are common across many cognitive tasks/domains (Assem et al., 2020; Fedorenko et al., 2013; Hodgson et al., 2021), which may include phonology (Hodgson et al., 2021; Poldrack et al., 1999), whereas the ventral LIFG contributes selectively to challenging semantic tasks (Gao et al., 2021; Whitney et al., 2011, 2012). One possible explanation is that ventral LIFG is specifically involved in controlled semantic retrieval processes as opposed to domain-general selection mechanisms, which are under the purview of dorsal LIFG regions (Badre & Wagner, 2007; Barredo et al., 2015; but see <u>Crescentini et al., 2010; Snyder et al., 2011)</u>. Alternatively, the processes implemented might be equivalent, but connectivity differences mean that they operate on distinct sets of inputs/outputs.

The anterior and posterior LIFG clusters were each affiliated with networks that occupy different positions along a macroscale cortical hierarchy that transitions from sensorimotor to transmodal cortex (Margulies et al., 2016). Specifically, the posterior LIFG was connected with the VAN and SMN, which process inputs from the external environment (Corbetta et al., 2008; Menon & Uddin, 2010). In contrast, anterior LIFG was preferentially coupled with regions of the FPN and DN, which are positioned towards the top end of the cortical hierarchy (Margulies et al., 2016). The anatomical and functional separation of anterior LIFG regions from sensorimotor systems might be requisite for the implementation of perceptually-decoupled, temporally-extended, and higher-order cognitive control (Fuster, 2001; Kiebel et al., 2008; Raut et al., 2020; Taylor et al., 2015). This interpretation is consistent with the proposal that the PFC is characterized by a posterior-anterior gradient of hierarchical control (for a review, see Badre & Desrochers, 2019), which was motivated by studies showing that anterior PFC is preferentially engaged by tasks that require generalization over an extended set of rules, integration of a larger number of dimensions and/or contexts sustained over longer periods of time (Badre & D’Esposito, 2007; Bahlmann et al., 2015; Koechlin et al., 2003; Nee & D’Esposito, 2016; for an investigation focused on the LIFG, see Koechlin & Jubault, 2006). In the language domain, it has been suggested that LIFG has a key role in the integration of linguistic subordinate elements into superordinate representational structures, and that this reflects a caudal-rostral functional gradient from phonological to syntactic to conceptual processing (Hagoort, 2005; Uddén & Bahlmann, 2012; also see Asano et al., 2021; Jeon & Friederici, 2015).

### 4.3. Concluding remarks

Our analyses revealed two main axes of organisation in LIFG function, in anterior-posterior and dorsal-ventral orientations, which is consistent with broader proposals concerning the whole PFC (Petrides, 2005). Moreover, our results suggest that functional differentiation across the LIFG occurs in a graded manner, and we were not able to find any clear evidence for discrete functional modules. Crucially, we replicated the principal gradients using two independent measures of FC, which suggests that our results are not dependent on idiosyncrasies of the datasets, and instead reflect stable, generalizable properties of LIFG organisation. The high degree of cross-modal similarity also suggests that a comparable LIFG functional organisation underpins divergent mental states. Future work is needed to directly probe the functional significance of these organisational dimensions and assess the compatibility of our findings at different spatial scales (e.g., cellular) and within other neuroimaging modalities (e.g., tractography) such that it is possible to arrive at an integrated account of the functional organisation of the LIFG.

## Supporting information

Diveicaetal_LIFGgradients_SupplementaryInformation

## Open science practices and data/code availability statement

We used open data available via the Human Connectome Project (https://www.humanconnectome.org/) and the NeuroQuery database (https://neuroquery.org/). The code used for data pre-processing and analysis relies on open source software (e.g., (Abraham et al., 2014; Salo et al., 2022) and can be accessed at: github.com/DiveicaV/LIFG_Gradients. The gradient and functional connectivity maps can be accessed and visualized via NeuroVault at: https://neurovault.org/collections/ETPEFCDV/. Due to its exploratory nature, the study was not pre-registered.

## Acknowledgements

We acknowledge the support of the Supercomputing Wales project, which is part-funded by the European Regional Development Fund (ERDF) via the Welsh Government, and we are grateful to Aaron Owen for his help with using the associated resources. We thank Julio A. Peraza for compiling and providing access to annotations of Neurosynth terms. Data were provided [in part] by the Human Connectome Project, WU-Minn Consortium (Principal Investigators: David Van Essen and Kamil Ugurbil; 1U54MH091657) funded by the 16 NIH Institutes and Centers that support the NIH Blueprint for Neuroscience Research; and by the McDonnell Center for Systems Neuroscience at Washington University.

## Funding

This work was supported by the Economic and Social Research Council (ESRC) Wales Doctoral Training Partnership in the form of a PhD studentship [ES/P00069X/1] awarded to VD and RJB (PhD student: VD). Support for ARL, MCR, and RS was provided by that National Institute of Mental Health (R01-MH09606) and the National Institute of Drug Abuse (R01-DA041353).

## CRediT author statement

**VD**: Conceptualization, Methodology, Formal Analysis, Visualization, Writing – original draft, Writing – review and editing; **MCR**: Methodology, Software, Formal Analysis; **TS**: Methodology, Software; **AL**: Methodology, Resources, Writing – review and editing; **RLJ**: Conceptualization, Methodology, Writing – review and editing; **RJB:** Conceptualization, Methodology, Writing – review and editing, Supervision.

## Footnotes

1 The supplementary materials can be accessed at: https://osf.io/u2834/.

## Notes

https://osf.io/u2834/

### Competing Interest Statement

The authors have declared no competing interest.

