## Supplementary material for "Graded functional organisation in the left inferior frontal gyrus: evidence from task-free and task-based functional connectivity": Diveicaetal_LIFGgradients_SupplementaryInformation

**for**

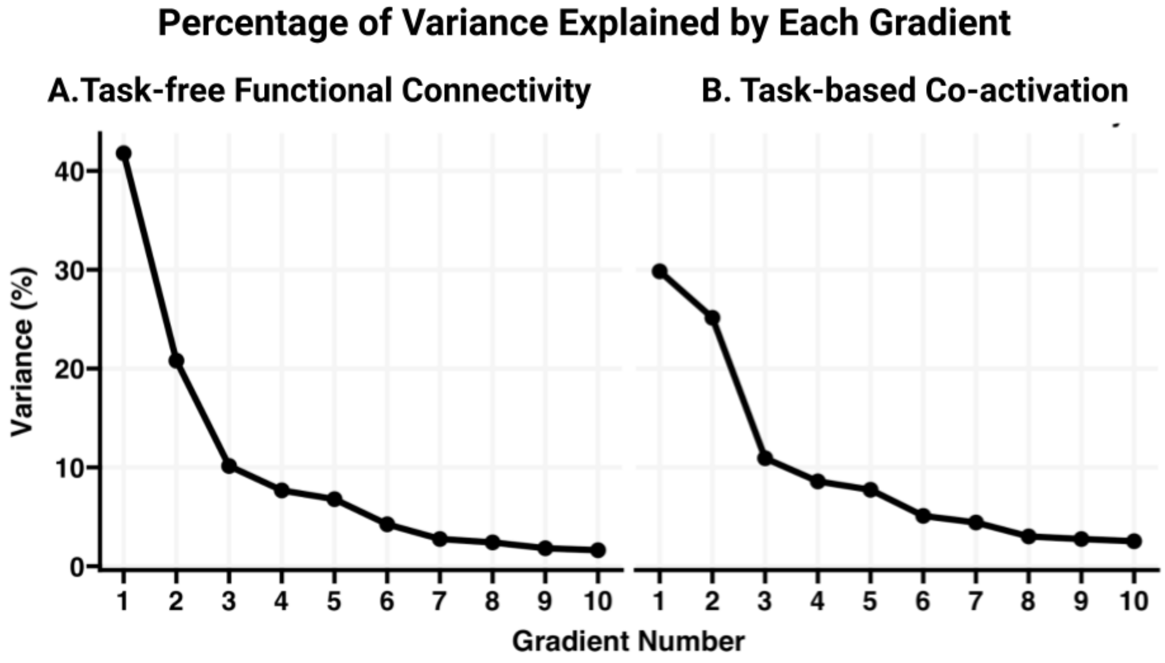


**Figure S1.** Percentage of variance explained by the 10 gradients derived from A) the task-free functional connectivity data and B) the task-based co-activation patterns.

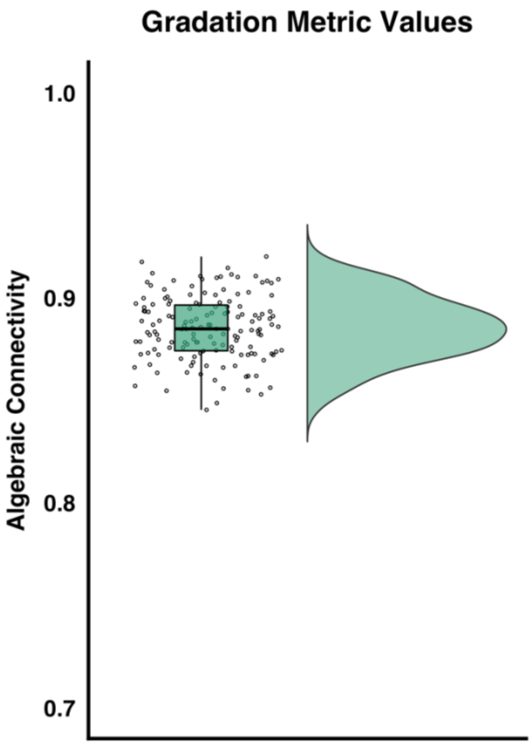


**Figure S2.** The distribution of the algebraic connectivity values (equivalent to the second largest eigenvalue of the Laplacian of the similarity matrix) obtained per participant in the task-free functional connectivity assessment are illustrated alongside individual datapoints and a boxplot highlighting the median, $25^{th}$ and $75^{th}$ quartiles. Values near 0 reflect the existence of hard clusters, whereas higher numbers suggest a graded change in functional connectivity. Note that the y-axis starts at 0.7, which is above the midpoint of possible values. The individual-level gradation metric values suggest that the left IFG is characterized by graded changes in task-free functional connectivity.

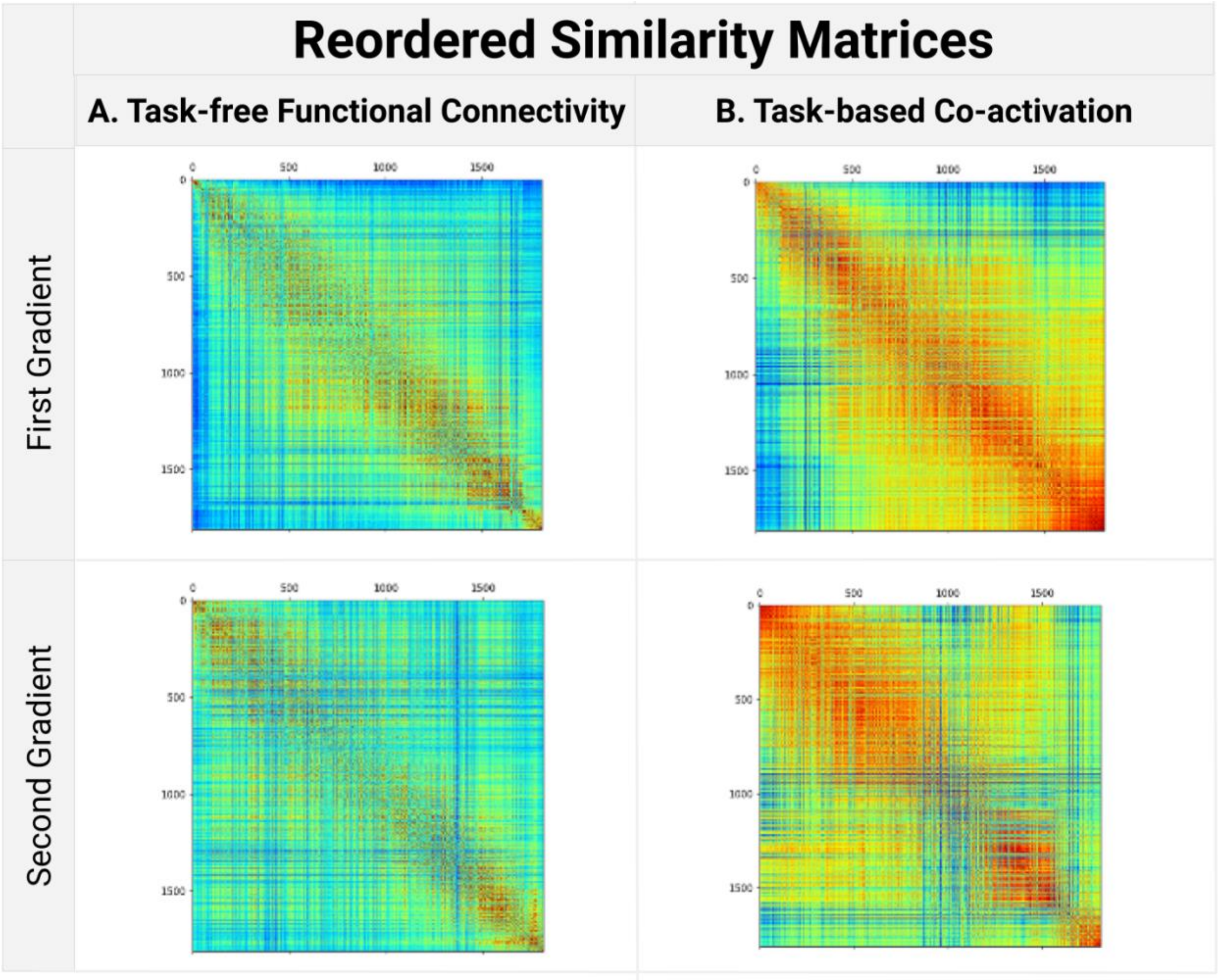


**Figure S3.** Similarity matrices reordered based on the voxels' positions along the first and second gradients. A) Reordered task-free FC group matrix. B) Reordered task-based co-activation matrix. Visual inspection of the reordered matrices suggests a high degree of gradation in the main axes of functional connectivity change across the left IFG.

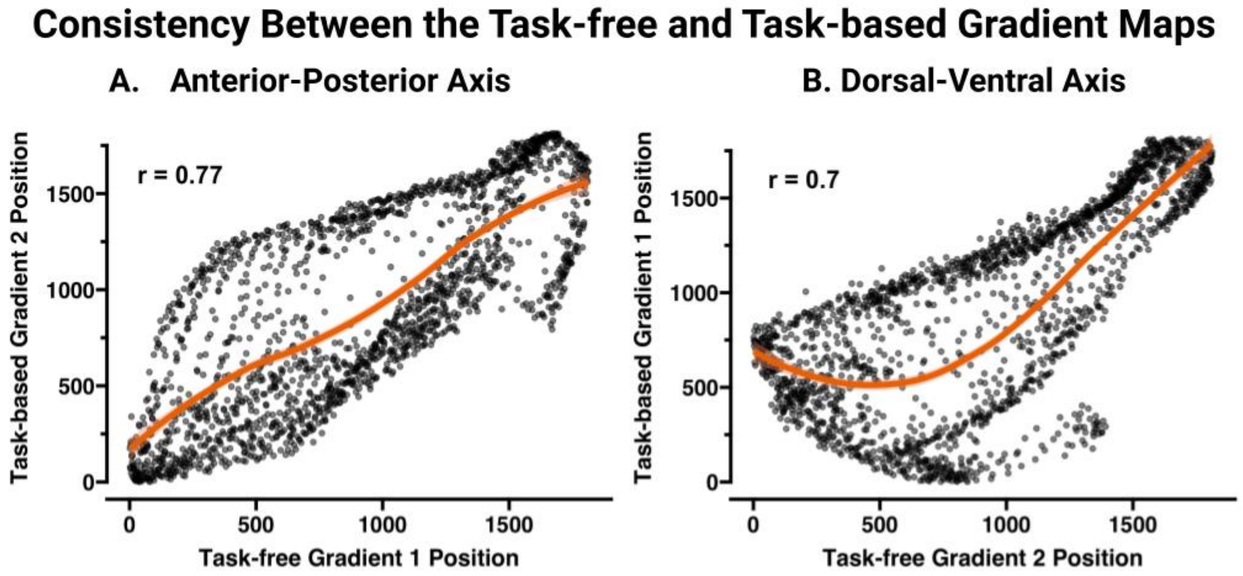


**Figure S4.** Scatterplots illustrate the relationship between voxels' positions in the resting-state gradients and the task-state gradients. The loess lines depicted in orange highlights the functional relationship. A. Voxels' ranks on the anterior-posterior (first) task-free gradient are plotted against their ranks on the anterior-posterior (second) task-based gradient. B. Voxels' ranks on the dorsal-ventral (second) task-free gradient are plotted against their ranks on the dorsal-ventral (first) task-based gradient. The r values represent the product-moment correlation coefficients and suggest strong relationships between the gradients extracted from independent FC datasets.

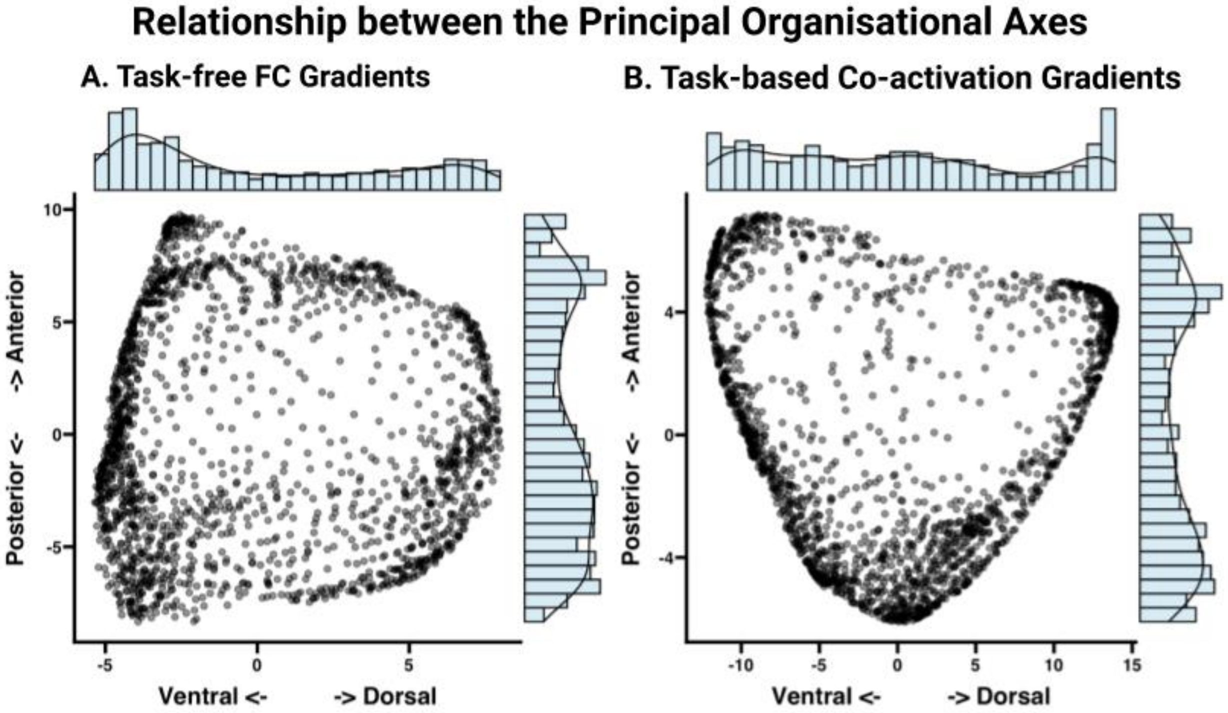


**Figure S5.** Scatterplots illustrate the relationship between voxels' gradient values on the first two connectivity embedding gradients extracted from A) task-free functional connectivity and B) task-based co-activation patterns. Histograms and density plots depicting the distribution of gradient values are presented on the respective axes.

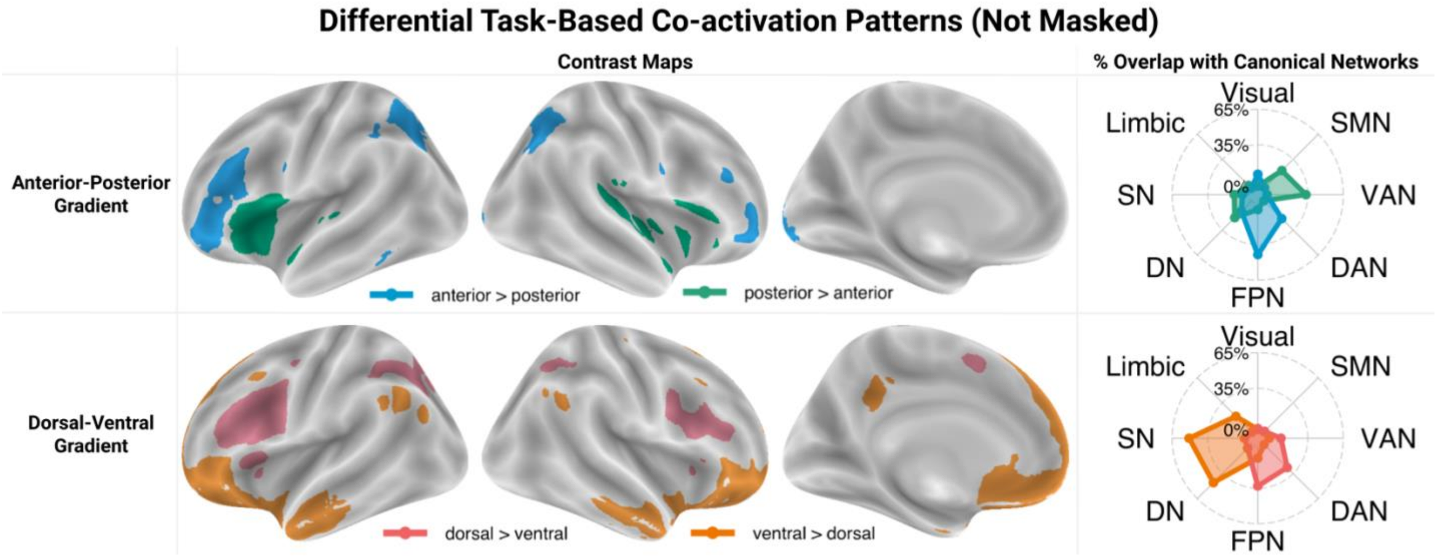


**Figure S6.** Results of contrast analyses between task-constrained co-activation patterns (derived using MACM analyses) of the IFG clusters located at the extremes of the anterior-posterior and dorsal-ventral task-based gradients. Unlike in the main text, these contrast maps were <u>not</u> masked using independent MACM maps. The spider plots in the right column show the percentage of overlap between the contrast maps and canonical networks from Yeo et al. (2011), as well as the semantic network from Jackson et al. (2016), which is comprised of regions that are functionally coupled with the ventrolateral anterior temporal lobe semantic hub at rest.

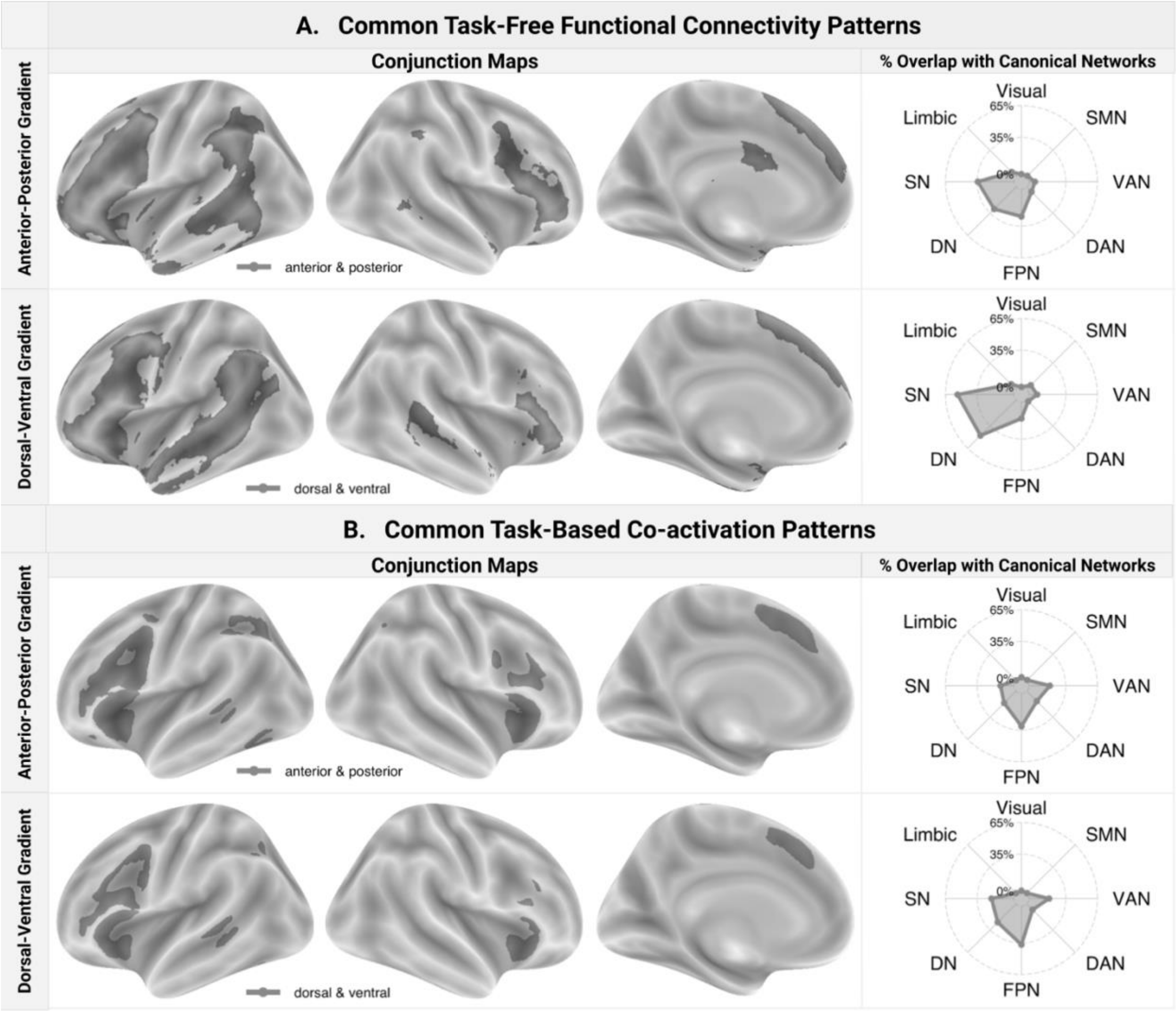


**Figure S7.** Conjunction maps showing common regions of (A) task-free functional connectivity and (B) co-activation between the hard clusters located at the extremes of the respective gradients. The spider plots in the right column show the percentage of overlap between the contrast maps and canonical networks.

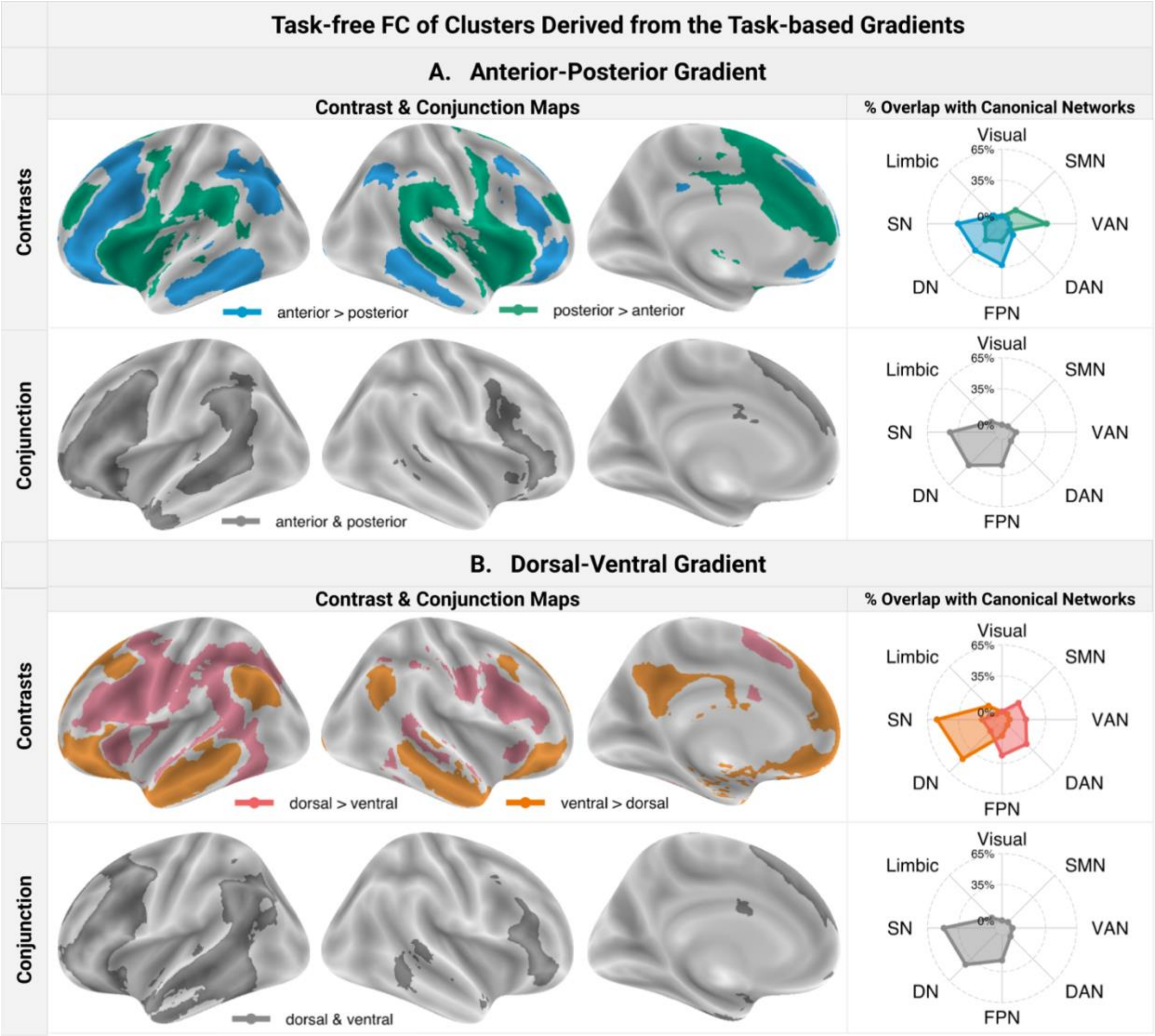


**Figure S8.** Contrast and conjunction maps showing regions of common and differential functionally coupling at rest between hard clusters representing the edges of the task-based FC (A) anterior-posterior gradient map and (B) dorsal-ventral gradient map. The spider plots in the right column show the percentage of overlap between the contrast maps and canonical networks.

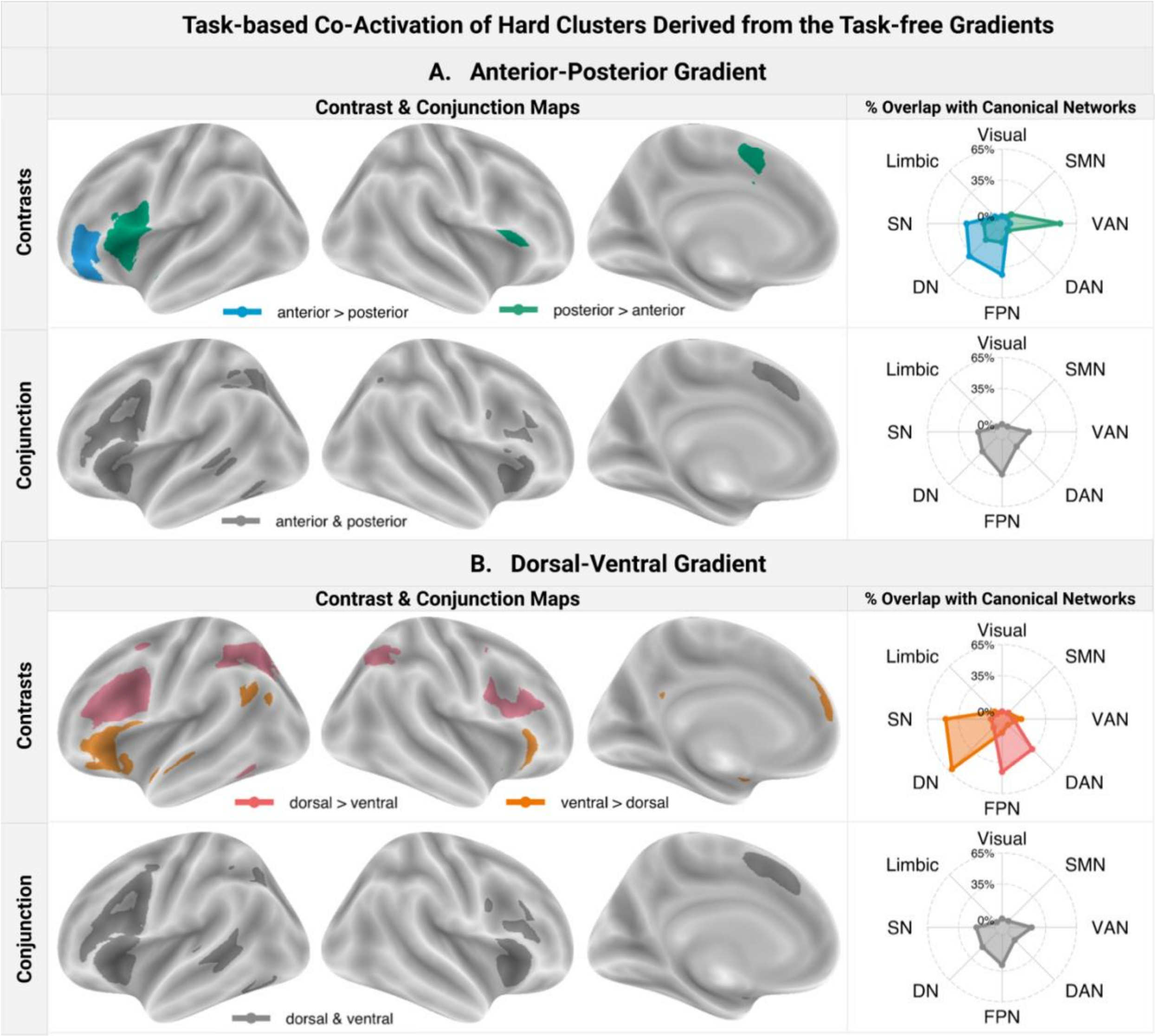


**Figure S9.** Contrast and conjunction maps showing regions of common and differential task-constrained co-activation across cognitive domains between hard clusters representing the edges of the task-free FC (A) anterior-posterior gradient map and (B) dorsal-ventral gradient map. The spider plots in the right column show the percentage of overlap between the contrast maps and canonical networks.

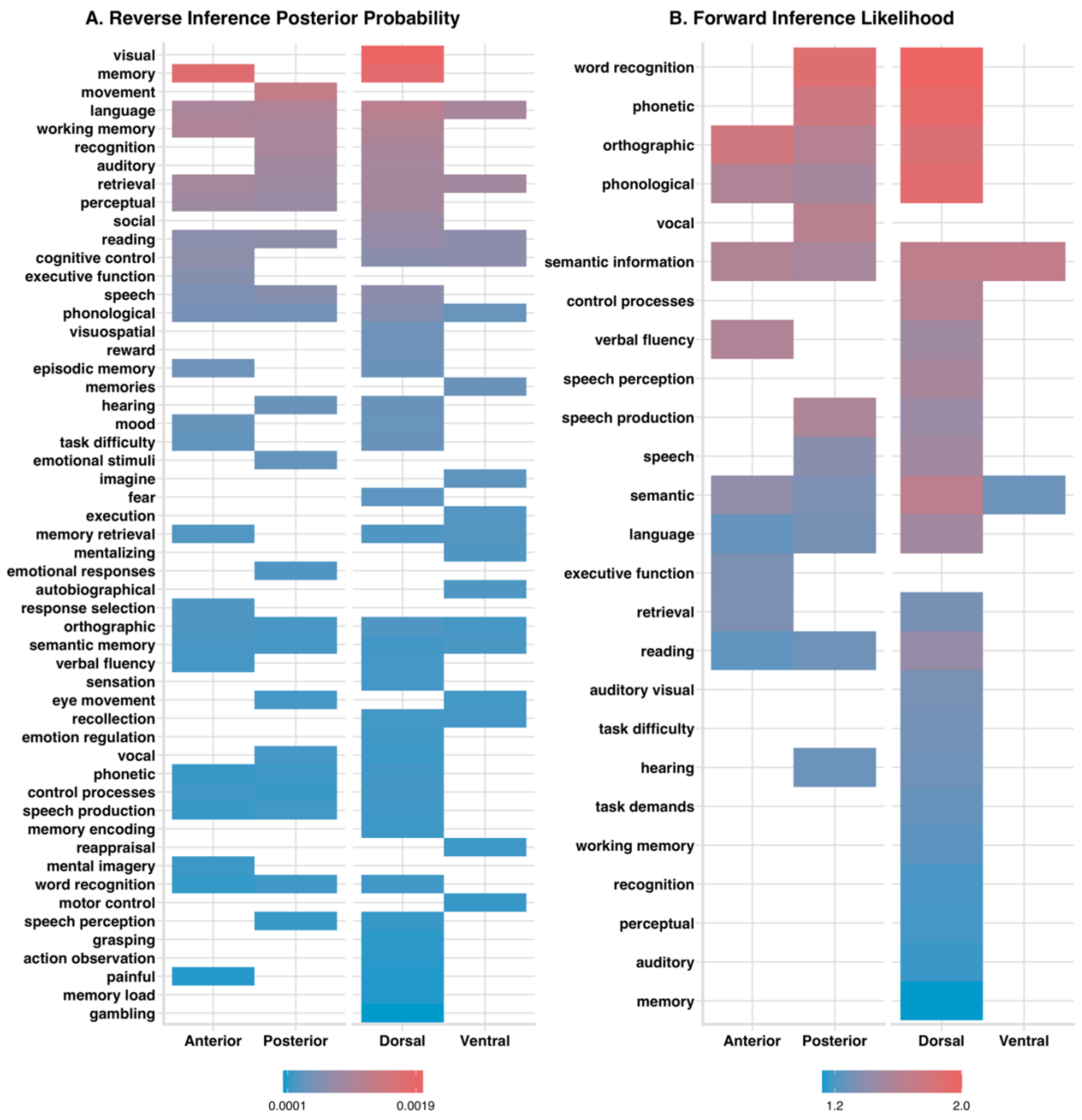


**Figure S10.** Functional terms associated with the IFG clusters derived based on the task-constrained gradients according to the A) specificity/reverse inference analyses and B) consistency/forward inference analyses. The colour indicates the effect sizes, with red colours suggesting greater association. Only statistically significant associations are highlighted. Synonymous terms with similar pattern of associations across the LIFG clusters were excluded.

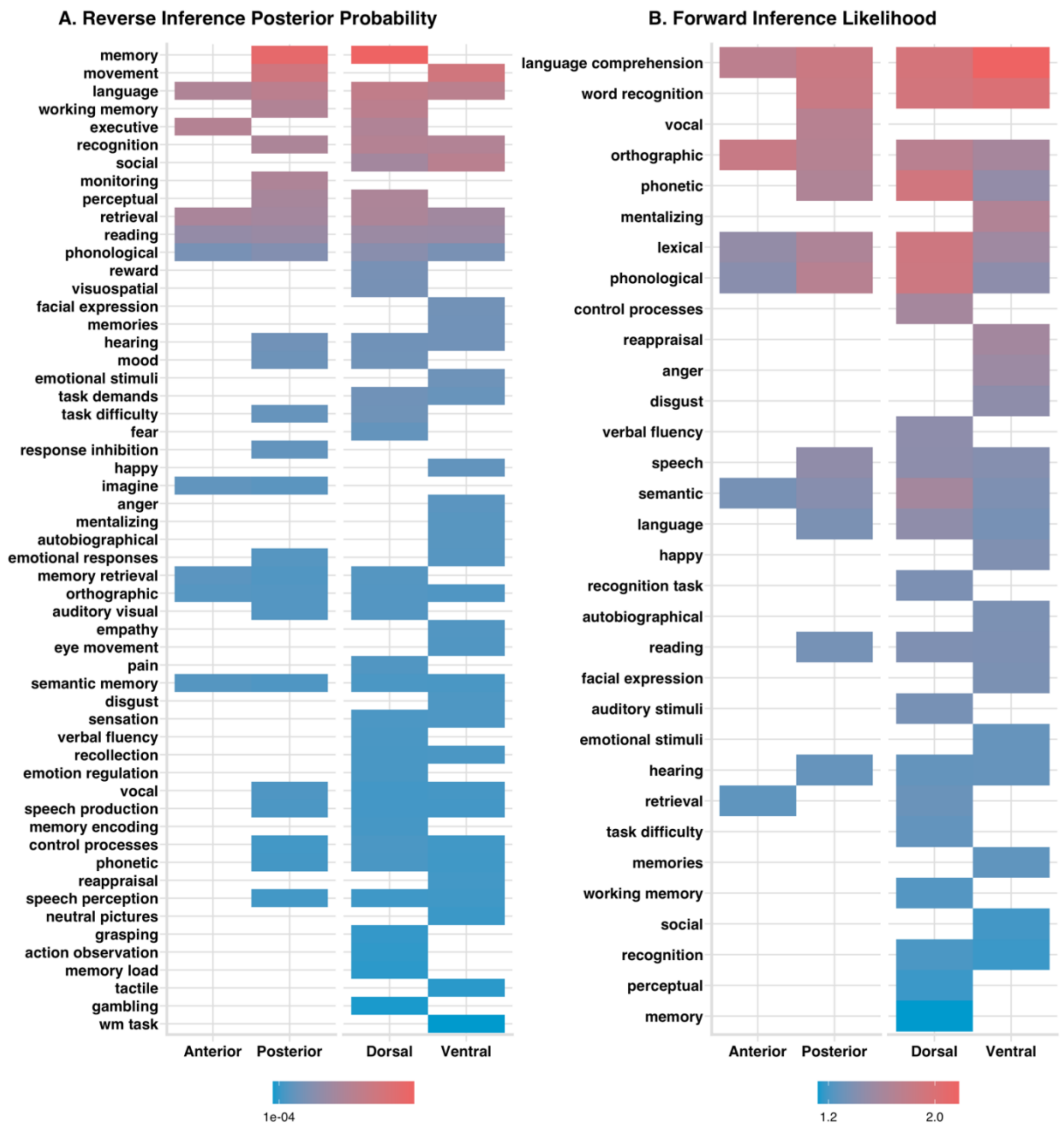


**Figure S11.** Functional terms associated with the IFG clusters derived based on the task-free gradients according to the A) specificity/reverse inference analyses and B) consistency/forward inference analyses. The colour indicates the effect sizes, with red colours suggesting greater association. Only statistically significant associations are highlighted. Synonymous terms with similar pattern of associations across the LIFG clusters were excluded.

**Table S1.** The MNI coordinates for the center of gravity of the hard clusters representing the edges of the task-free and task-based gradient maps.

| Cluster | Task-free Gradients | | | Task-based Gradients | | |
|---|---|---|---|---|---|---|
| | X | Y | Z | X | Y | Z |
| Anterior | -44 | 40 | -6 | -48 | 39 | 1 |
| Posterior | -50 | 17 | 9 | -44 | 21 | 2 |
| Dorsal | -50 | 23 | 21 | -51 | 21 | 22 |
| Ventral | -44 | 27 | -5 | -44 | 38 | -11 |

**Table S2.** The number of studies from the NeuroQuery database that reported at least one activation coordinate in each hard cluster. These studies were used as the input to MACM and functional decoding analyses.

| Cluster | Task-free Gradients | Task-based Gradients |
|---|---|---|
| Anterior | 664 | 851 |
| Posterior | 1064 | 1164 |
| Dorsal | 1332 | 1298 |
| Ventral | 1098 | 627 |

**Table S3.** Results of the seed-based resting-state functional connectivity analyses conducted on clusters extracted from the **anterior-posterior task-free gradient**.

| Analysis | AAL Label | Cluster Size (mm$^3$) | Max Z Value | X | Y | Z |
|---|---|---|---|---|---|---|
| Anterior Cluster > Posterior Cluster | Frontal_Inf_Orb_L | 51,200 | 21 | -36 | 38 | -12 |
| | Angular_L | 18,712 | 18 | -44 | -68 | 44 |
| | Frontal_Inf_Orb_R | 6,832 | 17 | 38 | 38 | -12 |
| | Cerebelum_Crus2_R | 17,824 | 16 | 46 | -72 | -40 |
| | Temporal_Mid_L | 16,336 | 16 | -64 | -44 | -10 |
| | Angular_R | 6,848 | 15 | 40 | -72 | 46 |
| | Cerebelum_Crus2_L | 10,336 | 14 | -42 | -72 | -40 |
| | Temporal_Mid_R | 6,568 | 14 | 62 | -40 | -8 |
| | Rectus_R | 4,216 | 14 | 4 | 44 | -16 |
| | Frontal_Inf_Tri_L | 912 | 13 | -52 | 24 | 28 |
| | Frontal_Mid_R | 2,952 | 11 | 32 | 20 | 50 |
| | Cerebelum_9_R | 520 | 11 | 2 | -56 | -50 |
| | Precentral_L | 672 | 11 | -46 | 8 | 36 |
| | Frontal_Inf_Tri_R | 2,320 | 9 | 52 | 34 | 22 |
| | Vermis_10 | 512 | 8 | 2 | -48 | -34 |
| | Temporal_Pole_Mid_L | 1,000 | 7 | -32 | 8 | -38 |
| Posterior Cluster > Anterior Cluster | Frontal_Inf_Oper_L | 17,720 | 20 | -52 | 16 | 2 |
| | Cingulum_Mid_L | 35,744 | 19 | -6 | 14 | 38 |
| | Frontal_Inf_Orb_R | 17,056 | 18 | 50 | 18 | -4 |
| | SupraMarginal_L | 21,256 | 16 | -56 | -40 | 26 |
| | SupraMarginal_R | 14,000 | 16 | 58 | -30 | 32 |
| | Precentral_R | 2,032 | 15 | 54 | 6 | 42 |
| | Frontal_Mid_L | 5,776 | 14 | -30 | 50 | 24 |
| | Frontal_Mid_R | 2,760 | 14 | 34 | 46 | 30 |
| Anterior Cluster ∩ Posterior Cluster | Frontal_Inf_Tri_L | 26,192 | 22 | -54 | 20 | 20 |
| | Frontal_Inf_Tri_L | 5,216 | 22 | -54 | 22 | 4 |
| | Supp_Motor_Area_L | 23,232 | 19 | -2 | 22 | 60 |
| | Parietal_Inf_L | 32,240 | 18 | -54 | -44 | 48 |
| | Frontal_Inf_Tri_R | 4,320 | 14 | 48 | 40 | 0 |
| | Frontal_Inf_Orb_R | 1,120 | 14 | 56 | 30 | -2 |
| | Frontal_Inf_Tri_R | 1,704 | 13 | 58 | 26 | 18 |
| | Cerebelum_Crus1_R | 832 | 11 | 14 | -76 | -30 |

| | | | | | |
|---|---|---|---|---|---|
| Temporal_Inf_L | 2,424 | 9 | -44 | -2 | -44 |
| Temporal_Inf_L | 856 | 8 | -50 | -6 | -40 |
| Fusiform_L | 608 | 8 | -44 | -40 | -18 |

**Table S4.** Results of the seed-based resting-state functional connectivity analyses conducted on clusters extracted from the **dorsal-ventral task-free gradient**.

| Analysis | AAL Label | Cluster Size (mm$^3$) | Max Z Value | X | Y | Z |
|---|---|---|---|---|---|---|
| Dorsal Cluster > Ventral Cluster | Frontal_Inf_Tri_L | 24,592 | 19 | -44 | 32 | 18 |
| | Parietal_Sup_L | 26,328 | 18 | -26 | -70 | 46 |
| | Precentral_L | 968 | 16 | -44 | 2 | 22 |
| | Temporal_Inf_L | 14,032 | 16 | -54 | -60 | -14 |
| | Frontal_Inf_Tri_R | 10,872 | 15 | 50 | 38 | 18 |
| | Frontal_Mid_L | 5,280 | 13 | -28 | 10 | 66 |
| | Temporal_Inf_R | 3,280 | 13 | 60 | -50 | -8 |
| | Cerebelum_8_R | 3,968 | 12 | 28 | -70 | -46 |
| | Cerebelum_Crus1_R | 832 | 10 | 6 | -80 | -24 |
| | Insula_L | 624 | 8 | -42 | -2 | 6 |
| Ventral Cluster > Dorsal Cluster | Frontal_Inf_Tri_L | 14,984 | 19 | -42 | 26 | 0 |
| | Frontal_Sup_Medial_L | 58,464 | 16 | -4 | 52 | 16 |
| | Insula_R | 12,960 | 14 | 32 | 20 | -14 |
| | Temporal_Inf_L | 13,808 | 14 | -48 | 2 | -36 |
| | Temporal_Mid_L | 10,728 | 13 | -58 | -18 | -10 |
| | Temporal_Inf_R | 9,960 | 13 | 48 | 2 | -32 |
| | Cerebelum_Crus1_R | 2,232 | 12 | 28 | -78 | -32 |
| | Temporal_Mid_R | 4,344 | 12 | 54 | -28 | -8 |
| | Angular_L | 12,208 | 11 | -54 | -60 | 30 |
| | Precuneus_L | 1,576 | 11 | -12 | -52 | 32 |
| | Temporal_Sup_R | 712 | 10 | 58 | -44 | 24 |
| | Cerebelum_Crus2_R | 648 | 9 | 24 | -88 | -38 |
| Dorsal Cluster ∩ Ventral Cluster | Frontal_Inf_Tri_L | 85,432 | 24 | -54 | 22 | 18 |
| | Frontal_Sup_Medial_L | 21,624 | 17 | -2 | 38 | 46 |
| | Frontal_Inf_Tri_R | 6,032 | 14 | 56 | 28 | 20 |
| | Cerebelum_Crus1_R | 7,016 | 14 | 14 | -80 | -30 |

| | | | | | | |
|---|---|---|---|---|---|---|
| | Temporal_Mid_R | 5,424 | 8 | 54 | -38 | 8 |
| | Fusiform_L | 5,272 | 8 | -30 | 2 | -44 |

**Table S5.** Results of the meta-analytic co-activation analyses conducted on clusters extracted from the **anterior-posterior task-based gradient**.

| Analysis | AAL Label | Cluster Size ($mm^3$) | Max Z Value | X | Y | Z |
|---|---|---|---|---|---|---|
| Anterior Cluster > Posterior Cluster | Frontal_Inf_Tri_R | 3,208 | NA | 48 | 36 | 18 |
| | Occipital_Mid_R | 5,432 | 4 | 34 | -68 | 34 |
| | Parietal_Inf_L | 9,816 | 4 | -38 | -44 | 42 |
| | Frontal_Inf_Tri_L | 20,392 | NA | -46 | 34 | 12 |
| Posterior Cluster > Anterior Cluster | Insula_R | 8,200 | NA | 44 | 18 | -2 |
| | Insula_L | 22,616 | NA | -42 | 18 | 0 |
| Anterior Cluster ∩ Posterior Cluster | Frontal_Inf_Oper_R | 18,200 | 4 | 44 | 20 | 10 |
| | Supp_Motor_Area_L | 15,224 | NA | -2 | 18 | 46 |
| | Pallidum_L | 1,648 | NA | -14 | 6 | 4 |
| | Parietal_Inf_L | 6,544 | NA | -36 | -54 | 44 |
| | Frontal_Inf_Oper_L | 41,856 | NA | -42 | 18 | 14 |
| | Occipital_Inf_L | 2,312 | NA | -44 | -60 | -12 |
| | Temporal_Mid_L | 1,208 | NA | -56 | -40 | 0 |

**Table S6.** Results of the meta-analytic co-activation analyses conducted on clusters extracted from the **dorsal-ventral task-based gradient**.

| Analysis | AAL Label | Cluster Size ($mm^3$) | Max Z Value | X | Y | Z |
|---|---|---|---|---|---|---|
| Dorsal Cluster > Ventral Cluster | Rolandic_Oper_R | 11,904 | 4 | 44 | 4 | 20 |
| | Angular_R | 2,408 | 4 | 32 | -56 | 48 |
| | Supp_Motor_Area_L | 4,976 | NA | -2 | 12 | 48 |
| | Parietal_Sup_L | 8,552 | 4 | -24 | -62 | 50 |
| | Frontal_Inf_Oper_L | 28,584 | NA | -46 | 16 | 22 |
| | Fusiform_L | 520 | 3 | -42 | -58 | -18 |

| | | | | | | |
|---|---|---|---|---|---|---|
| Ventral Cluster > Dorsal Cluster | Frontal_Inf_Orb_R | 4,048 | 4 | 36 | 22 | -18 |
| | Frontal_Sup_Medial_L | 1,080 | 4 | -6 | 42 | 40 |
| | Frontal_Inf_Orb_L | 17,152 | NA | -40 | 34 | -10 |
| | Angular_L | 2,720 | NA | -48 | -62 | 30 |
| Dorsal Cluster ∩ Ventral Cluster | Insula_R | 9,136 | NA | 40 | 22 | -4 |
| | Supp_Motor_Area_L | 9,800 | NA | -2 | 20 | 46 |
| | Frontal_Inf_Tri_L | 30,888 | NA | -44 | 20 | 12 |
| | Parietal_Inf_L | 2,696 | NA | -36 | -56 | 46 |
| | Temporal_Mid_L | 2,824 | NA | -56 | -38 | 0 |
